# Metagenome profiling and containment estimation through abundance-corrected k-mer sketching with sylph

**DOI:** 10.1101/2023.11.20.567879

**Authors:** Jim Shaw, Yun William Yu

## Abstract

Profiling metagenomes against databases allows for the detection and quantification of microbes, even at low abundances where assembly is not possible. We introduce sylph (https://github.com/bluenote-1577/sylph), a metagenome profiler that estimates genome-to-metagenome containment average nucleotide identity (ANI) through zero-inflated Poisson k-mer statistics, enabling ANI-based taxa detection. Sylph is the most accurate method on the CAMI2 marine dataset, and compared to Kraken2 for multi-sample profiling, sylph takes 10× less CPU time and uses 30× less memory. Sylph’s ANI estimates provide an orthogonal signal to abundance, enabling an ANI-based metagenome-wide association study for Parkinson’s disease (PD) against 289,232 genomes while confirming known butyrate-PD associations at the strain level. Sylph takes < 1 minute and 16 GB of RAM to profile against 85,205 prokaryotic and 2,917,521 viral genomes, detecting 30× more viral sequences in the human gut compared to RefSeq. Sylph offers precise, efficient profiling with accurate containment ANI estimation for even low-coverage genomes.

## 1 Introduction

Shotgun metagenomics, the sequencing of all microbes in a sample resulting in a *metagenome*, has allowed for unprecedented insight into microbial communities without the need for cultivating those communities in a wet lab [1,2]. Computational analyses often proceed by assembling metagenome-assembled genomes (MAGs) or profiling the reads against a database of reference genomes. While assembly is necessary for novel genome discovery, a fundamental drawback is that assembly does not work for low-abundance genomes. Reference-based profiling methods instead leverage vast collections of microbial genomes [3,4] to identify microbes and their abundances for even low-abundance organisms.

Metagenomes can be complex and large [5,6,7]. This necessitates accurate profiling methods that scale with high-depth samples and big databases. Algorithmic paradigms that have emerged under these constraints include efficiently finding short-exact matches from reads to a genome database [8,9,10] or sensitive read alignment against marker gene databases [11,12,13]. Short-exact match finding methods are known to have high numbers of false positives, so abundance cutoffs and confidence thresholds are now standard [14], especially for Kraken [8] and its derivatives [10]. Marker gene methods are more precise due to only retaining species-specific genes in their database, but such methods often use databases that are difficult to build and hard for users to customize.

An alternative algorithmic approach is k-mer *sketching* [15,16], where reads are subsampled via MinHash-derived [17,18,19] techniques into “bags of k-mers” called a sketch. This compressed representation allows quick average nucleotide identity (ANI) estimation of any reference genome against the genomes in a metagenome through containment k-mer statistics [20]. By contrast, popular methods such as Kraken or MetaPhlAn do not output nucleotide identity information. Although sketching methods tend to be efficient, previous implementations such as Mash screen [19] or sourmash [21] have accuracy issues for low-abundance genomes. This is due to k-mer content being missing [19] as a result of sequenced reads not fully covering the genome, obfuscating ANI calculation. Thus, arbitrary thresholds are also required to identify present genomes, an unsatisfactory solution given the importance of detecting low-abundance microbes [22].

### 1.1 Previous work

The issue of noisy sequence identity calculations due to low abundance has attracted attention in various other contexts. Genome-to-genome k-mer distance estimation [23,24] suffers from a similar issue when comparing two low-coverage single genome sequencing samples. In particular, our model uses similar assumptions as Skmer [23], but we deal with metagenomes instead of single genomes. Recently, ANI-based statistical testing of microbe presence-absence in metagenomes has been investigated in YACHT [25], but that requires a user supply a minimum coverage parameter in advance. Metagenomics gene presence-absence testing [26] is a related problem that deals with assembly incompleteness, but not necessarily low coverage. However, none of the aforementioned approaches deal with accurately estimating containment ANI in a metagenomics setting.

### 1.2 Our contribution

We present *sylph*, a k-mer sketching metagenomic profiler. The key innovation in sylph is a statistical model based on zero-inflated Poisson statistics to debias containment ANI under low coverage (**Fig. 1A**), solving the low-abundance ANI calculation problem. We show that sylph’s ANI estimation is accurate and apply it to species-level profiling through a principled 95% ANI cutoff [27].

**Fig. 1:**
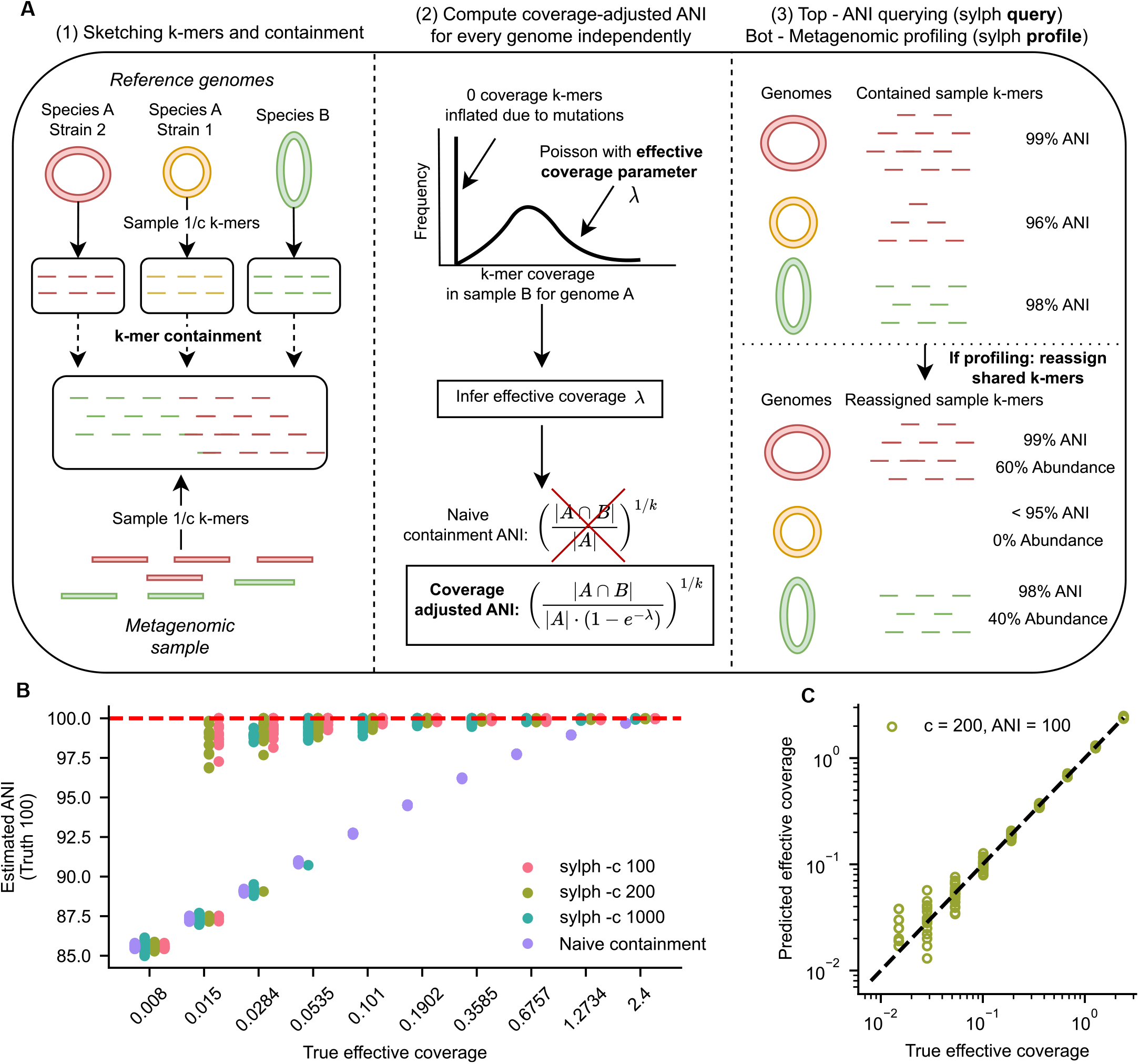
Algorithmic overview of sylph and demonstration for a synthetic dataset. **A**. (1) Sylph samples 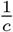 of the k-mers for metagenomes and reference genomes, resulting in *sketches*. Sylph computes the containment of the genome sketches within the sample sketch. (2) The coverage distribution of genome A’s k-mers within sample B is modeled with a zero-inflated Poisson distribution, where zero inflation is due to divergent k-mers between the genome and the metagenome having 0 coverage. Sylph infers the *effective coverage λ* and then estimates the final coverage-adjusted ANI. This is repeated for each genome within each sample. (3) Sylph can output the ANI estimates directly (sylph query) or a metagenomic profile (sylph profile), which reassigns shared k-mers and reports genomes with *>* 95% ANI. **B**. sylph’s ANI estimator over varying *c* (the subsampling rate) for a *K. pneumoniae* genome against simulated reads. Effective coverage is shown on the x-axis in log scale (20 samples per coverage). Naive indicates no coverage adjustment. **C**. Predicted and true effective coverage for the simulated *K. pnumoniae* sample with *c* = 200 (default) and 100% ANI between the sample and genome.

As a profiler, sylph is unique because it can estimate containment ANI *and* abundances, thus offering two orthogonal types of information. We display this versatility by finding ANI-based disease-strain associations on large metagenomic cohorts while also showing that sylph’s taxonomic profiling is both more precise and orders of magnitudes faster than existing profilers. Sylph is memory-efficient and allows for flexible database choice, enabling the usage of massive catalogs of genomes to improve the profiling of undercharacterized microbiomes.

## 2 Results

### 2.1 Accurate containment ANI estimation by zero-inflated Poisson k-mer statistics

Sylph first estimates the *containment* ANI [18] between a reference genome and the genomes in a metagenomic shotgun sample by searching the genome against the reads. A mathematical definition of containment ANI can be found in **Methods**, but intuitively, it measures the similarity of the reference genome to the genomes in the metagenome. Sylph first transforms a database of reference genomes and a metagenome into subsampled k-mers (31-mers by default) using FracMinHash [21], sampling approximately one out of *c* k-mers (*c* = 200 by default). Sylph then analyzes the containment [20] of the genomes’ k-mers in the metagenome (**Fig.1A, left**). While the containment ANI is known to be slightly biased upwards for *k* = 31 [19] (**Supplementary Fig. 1**) compared to standard ANI, this bias subsides for high ANI (*>* 95%). Therefore, we only work at the species level and refer to containment ANI as just ANI when the context is clear.

The key new methodological innovation in sylph is a statistical model to debias the containment computation for low coverage genomes. Under classical stochastic sequencing assumptions [28], we show that the distribution of a reference genome’s k-mer multiplicity in the metagenome can be modeled to follow a zero-inflated Poisson distribution with Poisson parameter *λ* and additional zero inflation. The zero inflation is due to base-level differences between the genome and the metagenome. We call the *λ* parameter in this zero-inflated Poisson the *effective coverage*; it takes into account k-mer lengths, read lengths, and read error rates. Sylph attempts to infer *λ* from the distribution and then correct for the missing k-mers to obtain a coverage-adjusted ANI estimate (**Fig. 1A, middle**).

After ANI estimation (**Fig. 1A, right**), sylph can output the ANI results directly using the sylph *query* command. However, this is not a *metagenomic profile* – a metagenomic profile includes the relative abundances of present taxa, but shared k-mers between similar genomes obfuscates which genomes are present and their abundance calculation. To resolve this, the sylph *profile* command reassigns shared k-mers to the highest ANI genome for each k-mer and recomputes coverage-adjusted ANI. Finally, we output the genomes with *>* 95% containment ANI and their abundances after reassignment while optionally estimating the percent of undetected sequences within a sample.

To test sylph’s ANI estimator under a baseline, we simulated error-free short reads using ART [29] for a single *Klebsiella pneumoniae* genome, generating single genome shotgun samples. This simulation shows that naive containment ANI (“naive ANI” for short), which sylph can also output, noticeably underestimates ANI when coverage is *<* 1 (**Fig. 1B**). However, beginning at 0.015 effective coverage, sylph can correct ANI to 95% species-level for some samples, owing to accurate estimation of the effective coverage *λ* (**Fig. 1C**). Additional experiments for when the reference genome has 96% and 90% containment ANI instead are shown in **Supplementary Figs. 2**,**3**. In general, the more similar a genome is to the reference genomes in the database, the better the coverage adjustment.

### 2.2 Sylph’s profiling is more precise and faster for the CAMI2 challenge datasets and divergent metagenomes

We first constructed an undercharacterized synthetic metagenome representing an environment where most of the genomes are not present in the database at the species level, but present at the genus level. Specifically, these synthetic communities had 50 present species at 95-97.5% ANI and 150 genomes between 85-90% ANI to the nearest ANI genome in the database (**Methods**). We compared sylph’s species-leveling profiling against three other methods, Bracken [10] (with Kraken2 [8]), KMCP [30], and ganon [9]. We chose Bracken due to its wide use, KMCP because it achieved the best CAMI2 marine benchmarks in its publication, and ganon because it is relatively time and memory-efficient. Importantly, we used the GTDB-R89 [31] database for all methods to standardize results. All methods were run with default parameters except Bracken, for which we used a 0.01% sequence abundance (as opposed to taxonomic abundance [32]) threshold.

On this undercharacterized synthetic metagenome (**Fig. 2A**), all methods except for sylph perform poorly for species-level classification, with precisions *<* 50% and F1 *<* 60%. On the other hand, sylph has 92% precision and 82% F1. This is likely due to sylph using a principled ANI cutoff; it does not mistake genus-level presence as species presence, therefore also having the lowest normalized L1 sequence abundance distance at the species level. Species-level abundances obtained by sylph can be aggregated to obtain abundances for higher ranks, but sylph does not attempt higher-level classification as part of it’s algorithm. Thus, when all genomes (including those only present at genus level) are included, sylph has a high L1 distance. Nevertheless, all profilers except for Bracken display a high L1 distance as well, indicating that KMCP and ganon likewise can only detect abundances accurately at the species level.

**Fig. 2:**
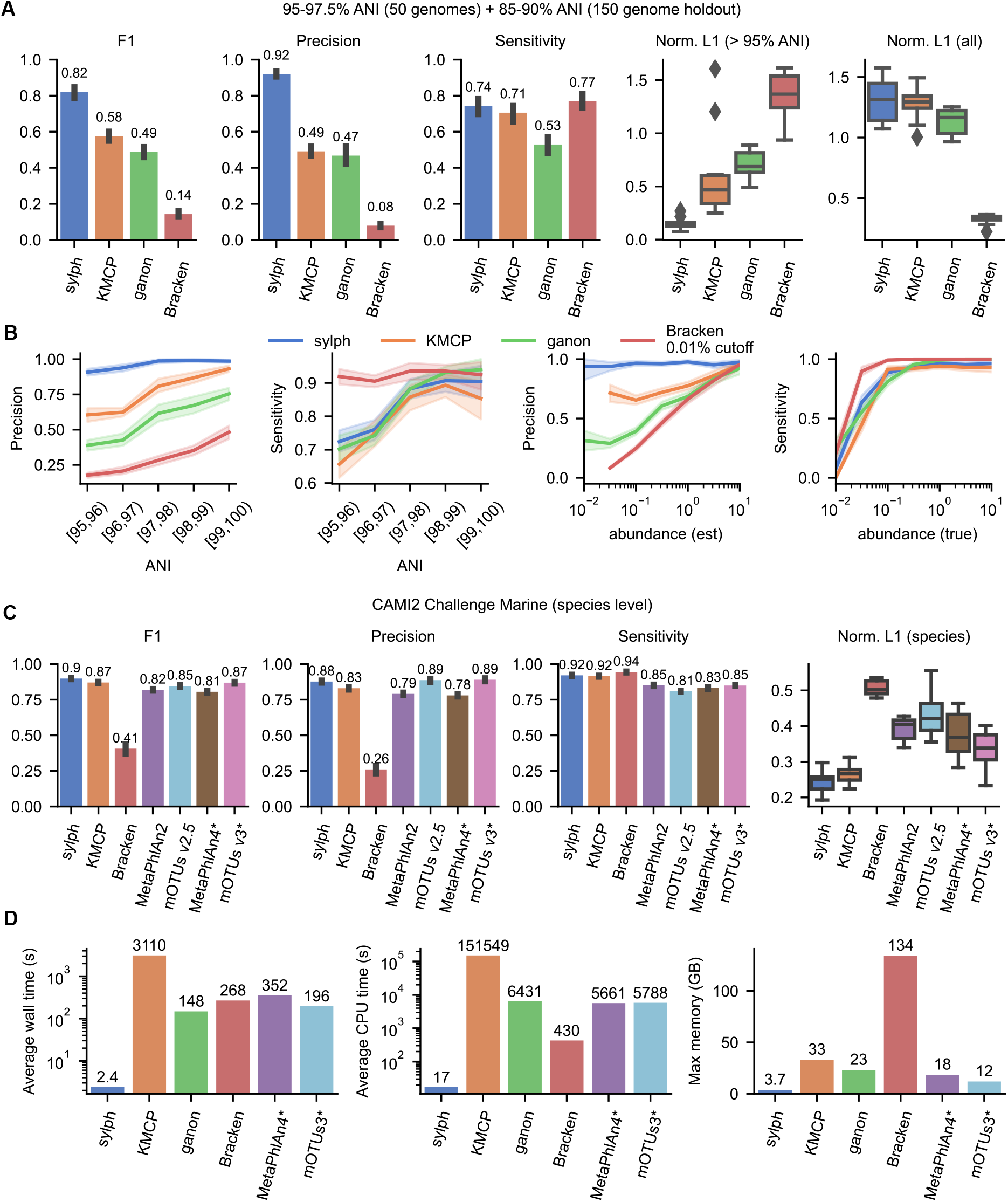
Profiling synthetic metagenomes with the GTDB-R89 database and the CAMI2 marine challenge. **A**. Species-level profiling for a community with 200 genomes: 50 species-level genomes with 95-97.5% ANI to the closest genome in the database and 150 genomes with 85-90% ANI to the closest genome (considered as not present at the species level). All methods used the GTDB-R89 database. The left boxplot’s L1 distances consider only the 50 species genomes, with all other abundances set to 0. The right boxplot normalizes L1 distances over all genomes. **B**. Five communities with 50 genomes stratified into 1% nearest neighbor ANI bins (10 samples per bin). All methods used the GTDB-R89 database. The rightmost two plots show the profiling precision/sensitivity over all ANIs (all 50 samples) as a function of sequence abundance binned by 0.5 in log space. The method’s estimated abundance is labeled “est” while the true sequence abundance is labeled “true”. **C**. Species-level results on the CAMI2 marine challenge data set. Methods with asterisks use different databases than the provided CAMI RefSeq database. **D**. Runtimes and max memory usage for the 200 genome dataset. MetaPhlAn4 and mOTUs3 were run on default databases, whereas other methods used GTDB-R89. 95% confidence intervals are shown in bar plots and line plots. Boxplots show the median, interquartile range (IQR), and 1.5 times the IQR.

To see the effect of genome divergence on profiling, we created 5 synthetic metagenomes, this time sampling genomes by 1% ANI intervals from 95-96% to 99-100% (**Fig. 2B, left**) to the nearest genome in the database. A striking result is that sylph’s precision is uniquely robust to the ANI between database and sample, maintaining *>* 90% precision over all ANI values. When we grouped all bins together for a uniform [95, 100) dataset (**Fig. 2B, right**), sylph was the only method that had consistently high precision over all sequence abundances. We attempted to benchmark against MetaPhlAn4 on this dataset by linking its taxa to the GTDB-R89 taxonomy (**Supplementary Fig.4**, **Methods**), which was used by the other methods. MetaPhlAn4’s precision stayed high across different regimes, like sylph, but its sensitivity and precision were consistently lower. We found that MetaPhlAn4’s species-genome-bin (SGB) definitions made 1-to-1 taxonomy associations difficult, lowering both its sensitivity and precision on this dataset.

To fairly benchmark against marker gene methods that utilize fixed taxonomies and databases, we leveraged existing CAMI2 challenge [33] profiler results. These submissions all had access to the same taxonomy and database information. We ran sylph with provided taxonomies and references against a set of CAMI2 submissions for the Marine dataset (**Fig. 2B, Methods**). We also bench-marked against the Strain Madness dataset and display genus level results in **Supplementary Figs**. 5-7. Notably, we ran updated versions of MetaPhlAn (v4) and mOTUs (v3) ourselves because existing submissions use outdated versions; however, the new versions use different databases and taxonomy than the official CAMI2 submissions, so the comparisons are not strictly fair (see **Supplementary Methods B**). For the species-level comparisons, sylph had the highest F1 score on both the Marine and Strain Madness datasets. Sylph had the lowest median L1 error for the Marine dataset and the second lowest for the Strain Madness dataset at the species level, being slightly higher than MetaPhlAn4 (0.16 vs 0.14). Overall, it appears that sylph is better or at least as good as marker gene methods at the species level, but like any other high-quality benchmarking standard, we caution against overinterpreting the CAMI2 results. It may be easy for profilers to inadvertently over-optimize for the gold standard profiles, and we discuss some pitfalls in **Methods**. For runtime and memory usage, we benchmarked all methods on the 200 genome undercharacterized dataset (**Fig. 2D**) using 50 threads. Sylph is *>* 50 times faster than the next fastest method, ganon, for wall time while taking *<* 4 GB of memory for *>* 25,000 genomes, which is 30 times less than Bracken (134 GB). Sylph is *>* 100 times faster in terms of CPU time compared to other methods except Bracken (with Kraken2). Kraken2 did not utilize all cores efficiently; it spent most of the time loading a 134 GB database into memory for each run. On the other hand, sylph is engineered for multi-sample processing and is able to sketch and profile many samples at once.

### 2.3 Sylph’s statistical model generalizes to real short and long-read datasets

Sylph’s ANI model is based on Poisson coverage statistics, but such models may not hold in real samples due to effects such as GC-bias [35]. To evaluate sylph on real reads, We analyzed the 87-genome MOCK2 community from Meslier et al. [34] sequenced with Illumina, PacBio HiFi (*>* 99% mean identity, **Supplementary Methods A**), and Oxford Nanopore (older chemistry, 90% mean identity), all downsampled to 1 Gbp (**Fig. 2**).

Sylph displays ANIs that are closer to the true 100% value than the other methods when querying against a database of true reference genomes (**Fig. 3A**). Notably, on the nanopore dataset, the median ANI for the naive ANI, Mash screen, and sourmash were all less than 95% due to sequencing errors lowering the k-mer containment. However, because sylph can correct for effective coverage, which also takes into account sequencing error, sylph still gives good estimates (*>* 99% median ANI). When using GTDB-R214 as sylph’s database, sylph’s estimated ANIs (from the *query* option, not from the *profile* option) agree with the containment ANI between the GTDB genome and its nearest neighbour mock genome (**Fig. 3B**). This shows that sylph’s coverage-adjusted ANIs can be thought of as a nearest neighbor ANI query against a reference genome for simpler communities without too much shared k-mer content.

**Fig. 3:**
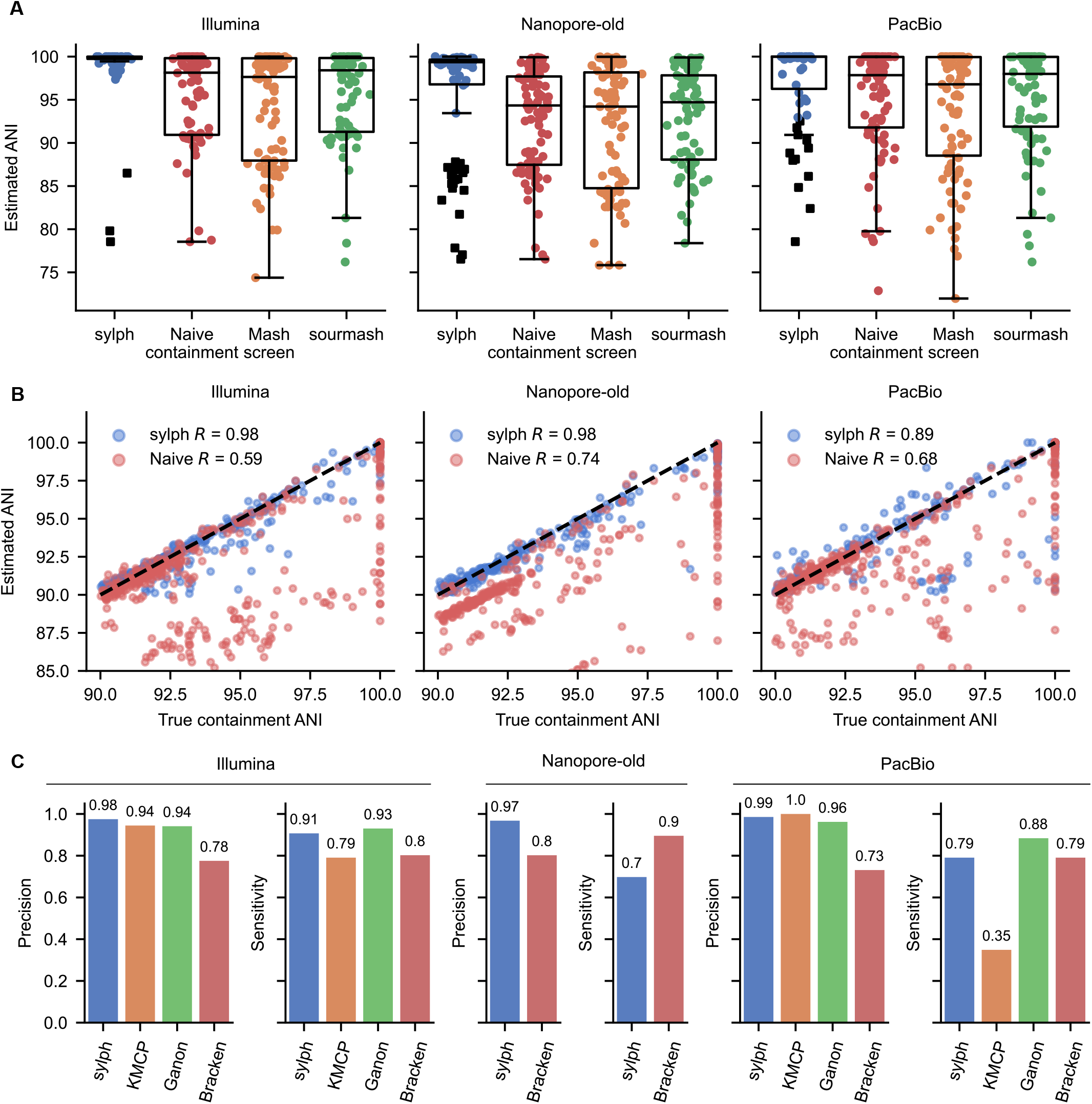
ANI and profiling results for real reads from a mock community [34] downsampled to 1 Gbp. **A**. Each method’s estimated ANI against the true reference genomes. Black squares indicate genomes whose coverage is too low for sylph’s coverage adjustment (but are still included in boxplot calculations). Nanopore-old denotes older chemistry nanopore reads with 90% sequence identity. **B**. Sylph’s coverage-adjusted ANI and naive ANI when querying against the entire GTDB-R214 database. The x-axis is the true containment ANI between a genome in GTDB-R214 against its nearest neighbor genome in the mock community. The y-axis is sylph’s estimated ANI. Pearson correlation coefficient values are shown. **C**. Species-level profiling results for the mock community. All methods used the GTDB-R89 database with default parameters except for Bracken, which used a 0.01% abundance cutoff. KMCP and ganon provided no output for the nanopore reads.

Using the same methods and databases for the previous profiling test, we profiled these three sets of reads and found that sylph was the most or second-most performant method on all datasets (**Fig. 3C**). The median ANI between the true reference genomes and its corresponding GTDB-R89 nearest neighbor genome was 100% for this dataset, making the profiling task much easier (as shown in **Fig. 2B**). Notably, sylph and Bracken were the only two methods that worked on the nanopore dataset by default, whereas the other methods output no results. Thus, sylph works on real reads with real biases, and in fact, it works on short, long, low-error, and even high-error reads. We attempted to benchmark this community with MetaPhlAn4 (**Supplementary Fig. 9**), but we found issues with its SGB taxonomy as in the previous benchmarking attempts. Regardless, MetaPhlAn4 only predicted 72 species *in total* on the Illumina dataset, whereas sylph and ganon predicted 80 and 81 *correct* species.

To benchmark sylph on real metagenomes without ground truths, we randomly selected 50 real short-read human gut metagenomes from GMrepo v2 [36] (accessions shown in **Supplementary Table 4**) and profiled them with sylph against the GTDB-R214 database. We then mapped the reads back to the genomes that sylph predicted were present and compared them against the coverage and abundance obtained by read mapping. While read mapping has its own biases for coverage and abundance calculation [37], we nevertheless use it as a simple ground truth. On this 50 metagenome dataset, sylph’s predictions are highly concordant with mapping-based abundance and coverage calculation (**Supplementary Figs 10** and **11**). The Pearson and Spearman correlation coefficients were (0.95, 0.95) for the abundances and (0.98, 0.95) for the coverages respectively, Sylph’s statistical model slightly overestimates the coverage for coverages *<* 0.1x (**Supplementary Fig 11**), likely due to slight bias of the estimator at very low coverages (**Fig. 1C**). We also compared sylph to MetaPhlAn4 (with identical taxonomies) and found that sylph’s number of predicted species was slightly higher but highly concordant (**Supplementary Fig. 12**) for species with *>* 0.01% abundance, although MetaPhlAn4 has a longer tail of species with ≤ 0.01% abundance. This is partially explained by sylph having higher abundance estimates than MetaPhlAn4 for low-abundance genomes, but given the lack of ground truth, we do not know if the long tail consists of false positives or false negatives.

### 2.4 A strain-level ANI-based metagenome wide association study (MWAS) for Parkinson’s Disease against 289,232 genomes

Sylph can estimate containment ANI between metagenomes and databases, giving genome similarity information. Because sylph’s containment ANI estimates work at low coverage, the ANI estimates work for even low-abundance genomes that *could not be assembled*. The containment ANI of every database genome in the metagenome obtained with the *query* option, can be viewed as a continuous presence (high ANI) and absence (low ANI) metric of a database’s genomes. Notably, this returns the ANIs at the genome level without collapsing at the species level.

A study by Wallen et al. [38] performed a gut MWAS for a cohort of 490 Parkinson’s Disease (PD) patients and 234 controls using relative abundance. Instead of relative abundance, we conduct a new MWAS using ANI as a covariate. As opposed to relative abundance, ANI is not compositional (**Discussion**), allowing for simpler analysis. Furthermore, relative abundance requires collapsing at a taxonomic level, whereas ANI does not. We use a simple logistic regression model for finding associations in two different ways: (1) with a continuous *>* 98% ANI covariate, which we use as the primary method and is shown in **Fig. 4, 4A**, and (2) with a 99% ANI threshold 0-1 variable. We searched for associations among all 289,232 genomes from the UHGG v2.01 catalogue [39] without dereplicating at the species level. Querying against all 289,232 genomes for 724 samples (5.5 tbps) took *<* 4 hours and 22 GB of RAM with 40 cores.

**Fig. 4:**
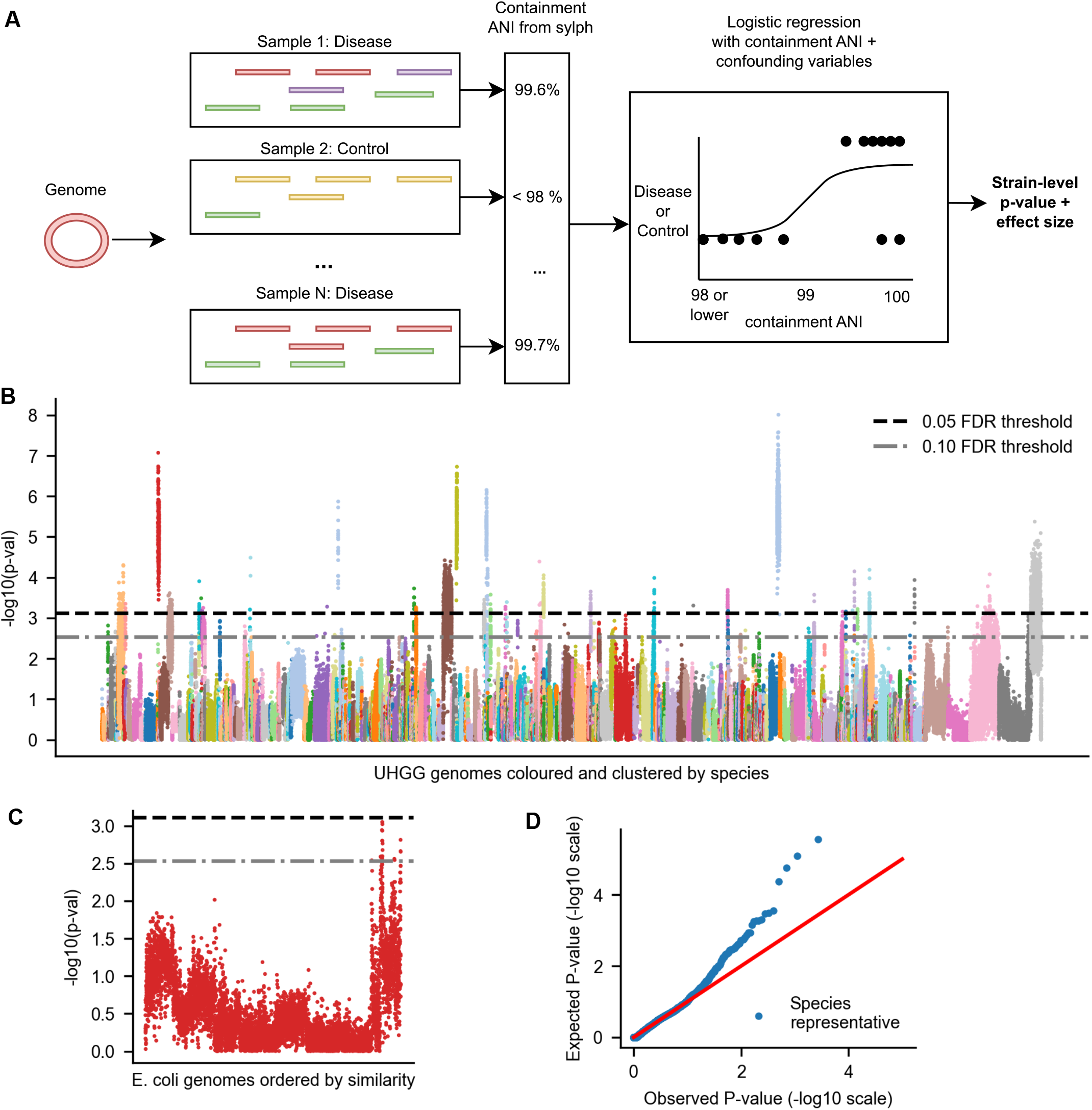
Sylph unveils strain-level ANI-disease associations for Parkinson’s disease (PD). **A**. Sylph associates a set of containment ANIs to query genome against the sample cohort. These containment ANIs are used as covariates for a logistic regression model. The p-values for this coefficient and the effect size are used for our associated study. **B**. Manhattan plot for an ANI-based MWAS of 289,232 genomes from UHGG against 724 Parkinson’s disease-control gut microbiomes [38] coloured by species. **C**. Q-Q plot where only one species representative was picked for each genome. **D**. A within-species linear ordering on the genomes was determined by clustering genomes by ANI similarity, unveiling strain-level variation for the p-values as shown for *E. coli*).

In Supplementary Table 3, we display the associations for genomes passing a 0.05 or 0.10 adjusted p-value threshold after multiple testing correction for methods (1) and (2) described above. 25 and 20 species have a genome passing the 0.05 threshold for methods (1) and(2) respectively, and the intersection had 13 species. The resulting Manhattan plot (only for the *>* 98% ANI method) is shown in **Fig. 4B**, along with its Q-Q plot in **Fig. 4D**. To order the genomes in the Manhattan plot, we created a linear ordering on the genomes within a species cluster based on genome similarity (**Methods**). This allows for visual inspection of the strain-level heterogeneity of p-values for *Escherichia coli* (**Fig. 4C**). Only 19/8309 *E. coli* genomes pass the 0.10 FDR threshold, and we found the passing genomes to be positively associated with PD, in agreement with Wallen et al. We searched the lowest p-value genome against a set *>* 4, 271 *E. coli* genomes downloaded from NCBI [27], and found a 99.99% ANI and 99% aligned fraction match with an *E. coli* isolate from a patient with bacteremia (PRJNA203078) as well as a 99.84% ANI match to a uropathogenic (UPEC) *E. coli* strain. Facultative anaerobes such as *E. coli* can be overrepresented in disease states but distinguishing between causality and correlation is difficult [40]. Nevertheless, the interpretability of genome-level results shown here provides an additional avenue for investigating correlations.

Of the 25 genomes passing the 0.05 FDR threshold for the 98% continuous ANI method, 5 of them were negatively associated with PD. Notably, three of the genomes were *Blautia wexlerae, Agathobacter rectalis* (also known as *Eubacterium rectale*), and *Roseburia intestinalis*, which are either producers or are associated with the production of short-chain fatty acids [41,42,43] including butyrate. The inverse butyrate-PD correlation is also supported by *Ruthenibacterium lactatiformans* associations, which is enriched in PD and contains the genome with the lowest p-value – the ratio of *B. wexlerae* to *R. lactatiformans* abundance was shown to be positively correlated with butyrate production [44] on a separate PD gut metagenome cohort and *R. lactatiformans* presence appears to be inversely correlated with butyrate production [45] in general. *Faecalibacterium prausnitzii*, a prominent butyrate producer [46], also had a significant genome at 0.10 FDR that was negatively correlated with PD but not 0.05 FDR. Ultimately, it appears that using ANI as the main covariate in MWAS leads to a simple but biologically consistent statistical analysis that *gives strain-level associations*, serving as candidates for further analyses, although we discuss power limitations in **Discussion**.

### 2.5 Sylph can integrate massive custom databases for characterizing understudied microbiomes, viruses, and eukaryotes

A key advantage of sylph over marker gene profiling methods is that sylph allows for the profiling of arbitrary genome databases as opposed to fixed databases. Sylph is technically a genome-level profiler and thus does not need taxonomy information, although taxonomy information can be added in post-processing. This allows for the profiling of prokaryotic MAGs that are not in public databases, as well as eukaryotes and viruses.

To show that sylph works on eukaryotic genomes, we analyzed skin metagenomes for an atopic dermatitis (AD) disease-control metagenome cohort from Chng et al. [47] with 19 AD patients and 15 controls. We investigated two prevalent fungal species, *Malassezia restricta* and *Malassezia globosa*. We used sylph to profile all samples against these two genomes and the GTDB-R214 database (accession in **Supplementary Table 2**). Of the detected *M. restricta* genomes, we found that 23% of them were below the 95% naive ANI threshold, whereas 63% of *M. globosa* genomes were below the 95% threshold (**Fig. 5A**), indicating the usefulness and necessity of coverage adjustment for low-coverage eukaryotic species. Like Chng et al., we found that *M. globosa* was differentially abundant between case and control (p = 0.0066, Wilcoxon rank-sum test) but not for *M. restricta* (**Fig. 5B**), highlighting the concordance of sylph with other approaches for eukaryote profiling.

**Fig. 5:**
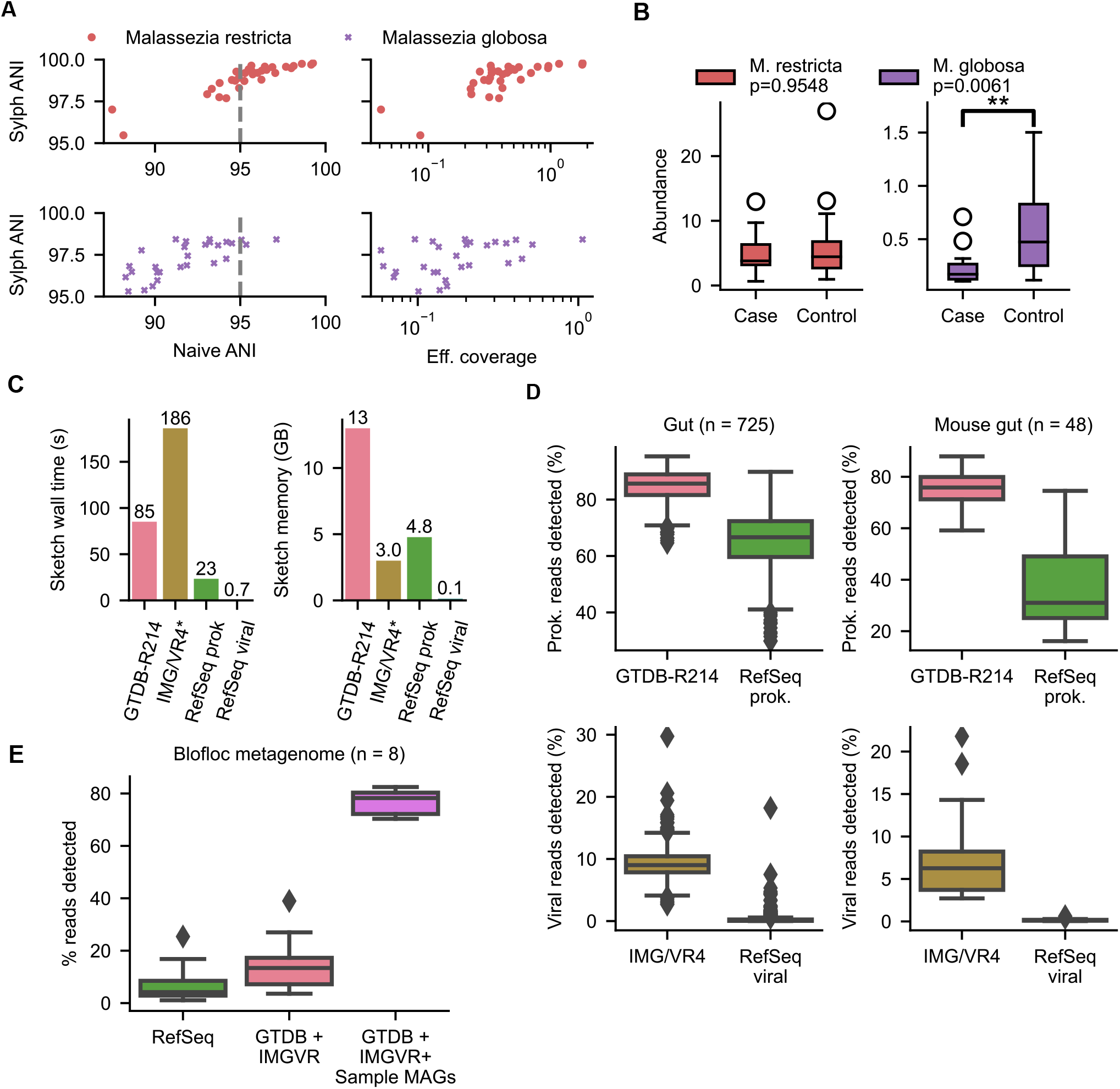
sylph can improve profiling completeness by incorporating eukaryotes, viruses, and arbitrary dereplicated databases. **A**. Sylph’s ANI for two fungal species against skin samples from Chng et al. [47] as a function of estimated effective coverage and naive ANI. **B**. Differential abundance analysis of the two fungal species between case and control AD patients calculated by the Wilcoxon rank-sum test gives concordant results to the previous study. *p *<* 0.05, **p *<* 0.01, ***p *<* 0.001. **C**. Wall times and max memory usage for sketching databases with 50 threads. **D**. Profiling with RefSeq representative genomes versus GTDB-R214 [31] or IMG/VR4[54] on human and mouse gut metagenomes. The read detection percentage indicates the number of reads that are accounted for by the genomes in the database detected at the species level. **E**. 8 biofloc metagenomes [50] are not well characterized by existing databases, but adding in their assembled MAGs increases the percentage of detected sequences.

To show sylph can profile viruses precisely, we simulated a viral metagenome from RefSeq and profiled against the MGV database [48] (**Methods, Supplementary Fig. 13**). We found that our coverage-adjustment procedure was less sensitive for viruses due to their smaller genome length, but the precision for profiling remains high even for smaller genomes. We then leveraged the IMG/VR4 database, a massive database of viral genomes clustered into 2,917,516 high-confidence operational taxonomic units, and the latest GTDB-R214 database with 85,205 prokaryotic genomes for profiling. Notably, these massive databases only took *<* 300 seconds and 16 GB of memory to index (**Fig. 5C**) with 50 threads, implying that sylph is efficient enough to leverage future database growth.

We profiled the Wallen et al. gut samples and also a mouse gut metagenome project [49] (PR-JNA549182) with the GTDB and IMG/VR databases, as well as with a database of representative RefSeq prokaryotic and viral genomes (**Fig. 5D**). Using GTDB-R214 and leveraging sylph’s ability to estimate how many reads are contained within the detected database genomes (**Methods**), sylph detected on average 84.8% human and 75.1% mouse reads belonging to prokaryotic species, whereas using RefSeq detected 75.1% human and 37.2% gut reads, undercharacterizing the mouse gut metagenomes at the species-level. The effect of virome profiling for RefSeq versus IMG/VR4 was much more pronounced, with 9.2% human and 6.9% mouse gut reads being detected as viral on average with IMG/VR4 versus only 0.3% human and 0.2% mouse gut with RefSeq.

While many microbiomes are still understudied and are not explained well by databases, sylph allows for customization of databases with MAGs to increase detection power. As an example, we took 8 recently published biofloc metagenomes [50] and found that existing databases did not adequately profile the metagenomes at the species level (**Fig. 5E**). By adding in the 444 MAGs assembled from these 8 samples, we found a drastic increase in detected reads, from 15.5% to 76.8% on average. Sylph’s coverage estimates were highly concordant with BWA [51] (**Supplementary Fig. 14**) on these novel MAGs (Pearson R = 0.995), and after indexing, sylph’s profiling was *>* 1000*×* faster than BWA’s mapping. Thus sylph can profile and calculate coverages against novel databases with MAGs in a manner concordant with read mapping. In **Supplementary Fig. 15**, we further show that adding in new catalogs of ocean MAGs and soil MAGs [52,53] can modestly improve the average percentage of detected reads for recently available samples of plant (25.5% to 36.1%) and ocean metagenomes (17.4% to 22.8%). Nevertheless, database incompleteness for species-level profiling is still an issue for understudied microbiomes (**Discussion**).

## 3 Discussion

sylph is a new method that uses k-mer sketching and zero-inflated Poisson statistics for containment ANI computation. When adapted into a metagenome profiler, it is more precise and orders of magnitude faster than other methods, especially for multi-sample profiling. Due to the ANI approach sylph uses for profiling, it has the unique ability to provide ANI estimates for metagenomes, even at low coverages.

The containment ANIs that sylph outputs, providing a continuous measure of presence (high ANI) and absence (low ANI), are in addition to its profiling abilities. We have shown that these ANIs are informant enough to give concordant results to previous PD studies based on relative abundance, yet containment ANIs offer genome-centric information at the strain level. Traditionally, relative abundances are used to predict functional implications for a microbiome, as is done in pathway analysis [55] or differential abundance analysis [56]. However, relative abundances are compositional [57,58] and are tricky to analyze due to spurious correlations [59] arising in compositional data. In contrast, containment ANIs are less compositional. An increase in a species’ relative abundance necessarily leads to the lower relative abundance of an unrelated, static species. However, changes in relative proportions should not affect the ANIs since the latter are based on genomic content. We believe that this approach and associated statistical techniques for downstream analysis deserve further exploration. A notable direction to explore is how to handle the loss of power caused by multiple testing correction in the presence of correlated strains for each species, analogous to the linkage disequilibrium problem in standard GWAS [60].

While sylph gives confident species-level predictions and accurate ANIs, further work is needed on fast and precise methods for profiling at higher taxonomic ranks. While sylph can explain much of the metagenomic reads in microbiomes such as the human gut or mouse gut, many environments can still not be classified well at the species level [11] due to database incompleteness. Our strategy for tackling this problem was to design sylph so that researchers can create customized databases from their novel genomes or MAGs, although this requires the generation of new genomes for researchers working in undercharacterized microbiomes. Despite the continuing efforts to build larger databases, this will be an issue for the foreseeable future given the small fraction of species-level diversity explained by high-quality assembled genomes.

## 4 Methods

### 4.1 ketched k-mer representation

We represent our sequences and genomes in terms of k-mers. We do not use all available k-mers in our sequences. Instead, we use a *sketch* [15] of all k-mers by using the FracMinHash method [21]: given a hash function *h* which maps k-mers to [0, *M*], we retain a k-mer denoted as *x* only if *h*(*x*) *< M/c*. The expected fraction of subsampled k-mers is exactly 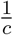 under appropriate uniformity and stochasticity assumptions on the hash function, so *c* can be thought of as the rate of subsampling. Our sketched representation has several benefits. Firstly, we take *c* = 200 by default, effectively speeding up computation and lowering memory costs by a factor of 200. Secondly, two subsampled k-mers are probably not near each other in a read or a genome, making them more independent of one another in a statistical sense, discussed later on.

### 4.2 Independent substitution model with read-sampling

Sylph assumes a generative probabilistic model, where calculating ANI becomes a question of parameter estimation under our model. The model consists of two parts: sequence evolution and read-sampling. We will consider the independent substitution model without spurious k-mers [61] and uniform random read-sampling with error.

Assume we are given a genome *S*. The first part of our random model is an independent substitution model on *S* parameterized by *θ* ∈ [0, 1]: assume every letter in *S* mutates to another letter with probability *θ* independently, resulting in a mutated version denoted *S*^*′*^.

#### Definition 1.

*Given random substitution parameter θ, let τ* = 1 − *θ. In our model, τ is defined as the (fractional) average nucleotide identity (ANI)*.

Note that we define ANI to be fractional for mathematical convenience.

The second part is a read-sampling step, independent of the first part, where reads are sampled from the new genome *S*^*′*^ uniformly at random. Each read has some error rate *ϵ*, where errors are also modeled as independent substitutions. Importantly, under the uniformity assumption and that the reads are small enough, we can approximate the coverage as Poisson distributed: if the average number of times a base is covered (its *true coverage*) is *δ*, the distribution of the number of times a base is covered follows a Poisson distribution with mean *δ*.

We will extend this Poisson assumption to k-mers. That is, the k-mer *multiplicity*, the number of times a specific k-mer is seen, is also Poisson distributed. However, due to the read errors, where every read has its own error rate *ϵ* subject to some distribution *D*_*ϵ*_, the actual coverage of k-mers will be modified to 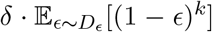, where a k-mer may not be “covered” due to it having an error within the read (note that errors are distinct from mutations between genomes). In addition, for a read of length *L*, there are only *L* − *k* + 1 k-mers, so the coverage decreases by a factor of (*L* − *k* + 1)*/L*. Assuming all reads have length *L*, we make the following definition.

#### Definition 2.

*Assuming read length L, k-mer length k, coverage δ, and read-error rate ϵ* ∼ *D*_*ϵ*_ *under a distribution D*_*ϵ*_, *we define the effective coverage λ as*

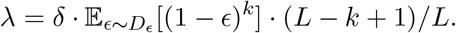

In practice, we observe k-mer multiplicities and not the true coverage, so the distribution is Poisson with parameter *λ*. That is, sylph only sees the effective coverage under our model and *not* the true coverage *δ*. We discuss true coverage computation in the section **Computation of effective and true coverage**.

### 4.3 Metagenome k-mer model

Under the above model, we can now state precisely the input and output for sylph. Let *R* be a set of reads from our sample. We model *R* as the output from uniform read sampling for a true underlying set of genomes *G*_1_, *G*_2_, … at effective coverages *λ*_1_, *λ*_2_, …, with all final reads mixed together. We will model an input genome 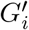 as being mutated from *G*_*i*_ due to random substitutions at ANI *τ*_*i*_. Our goal is to infer and output the true *λ*_*i*_ and *τ*_*i*_ parameters from our k-mer data under this model.

#### Definition 3.

*Given an input reference genome* 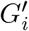 *and a fixed hash function resulting in N FracMinHash k-mers for* 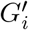, *let X*_1_, …, *X*_*N*_ *be the multiplicity of* 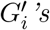 *k-mers in the reads R*.

We will make a few assumptions for modeling purposes. (1) The k-mers for each genome within the metagenome are unique. (2) If a k-mer is mutated, it does not appear in the metagenome. (3) If a k-mer in a read is erroneous, it does not show up in any genome. In practice, (1) is frequently violated due to mobile elements, which we take into account later on in the inference step, whereas (2) and (3) are assumed to be reasonable assumptions.

Under these assumptions, we derive our model of *X*_*i*_’s distribution as follows. If the k-mer for *X*_*i*_ is in *R*, then *X*_*i*_ is Poisson with parameter *λ*_*i*_. However, if the k-mer corresponding to *X*_*i*_ is mutated and thus differs between *G*_*i*_ and 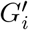, we assume it will not exist in *R*, so *X*_*i*_ is 0. The probability of a k-mer being not-mutated is 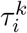, so it follows that

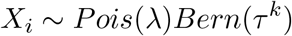

where the Poisson and Bernoulli random variables are independent by assumption. In other words, 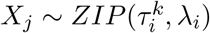 where *ZIP* is a zero-inflated Poisson distribution.

If the sampling locations of the FracMinHash k-mers for *X*_*i*_, *X*_*i*+1_, … are close together on the genome, these random variables become strongly correlated; nearby k-mers will be covered by the same reads frequently. However, we subsample k-mers by FracMinHash so on average our k-mers are *c* bases apart, which allows us to reasonably assume that our k-mers *X*_1_, …, *X*_*N*_ are sufficiently spread out so that independence of the *X*_*i*_s is not a bad assumption in practice.

### 4.4 Inference for *λ*

We can now rephrase our problem precisely as inferring the parameters *τ*^*k*^ and *λ* from *N* independent and identically distributed (i.i.d) samples *X*_1_, …, *X*_*N*_ ∼ *ZIP* (*τ*^*k*^, *λ*). There exists literature for inference concerning i.i.d ZIP samples, but existing MLE and MME estimators depend on means and variances of the *X*_*i*_s [62], which are not robust to long-tailed outliers that are commonly present in metagenomic data (e.g. mobile elements, low-complexity k-mers). As such, we use a ratio-of-multiplicity estimator for *λ* as used by Skmer [23], which we define below.

We first define *N*_*a*_ to be the number of *X*_*i*_s with multiplicity *a*. It follows that for all *a* ≥ 1,

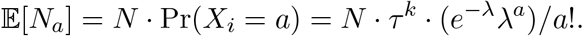

Notice that 𝔼 [*N*_*a*+1_]*/* 𝔼 [*N*_*a*_] = *λ/*(*a* + 1), which does not depend on *τ* . We have samples of *N*_*a*_ available from data, so we can use this ratio as our estimator.

#### Definition 4

**(Ratio-of-multiplicity** *λ* **estimator (inspired by** *ξ* **from Sarmashghi et al. [23]))**. *Our estimator for λ is*

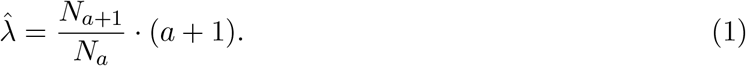

The estimator used in Skmer [23] differs from sylph in that their *N*_*a*_ variables are absolute k-mer frequencies over all k-mers in the sample, whereas our *N*_*a*_ variables are only k-mers contained in a reference genome.

In practice, the estimator has has reasonable bias except for very small *λ* (**Supplementary Fig. 3**). We discuss the zero denominator case in **Practical thresholds**.

### 4.5 *λ*-adjusted containment index and ANI

We described how sylph obtains *λ* in the previous section, but not the ANI *τ* . The central idea for calculating ANI is simple: check how many k-mers from the reference genome are contained in our reads. Let *A* be the set of sketched k-mers in our genome, and *B* be the set of sketched k-mers in our reads. We call 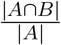 the *containment index* [20]. If *A* and *B* are k-mers from two complete genomes, without read-sampling, then the containment index can be transformed into an estimator for *τ* = *ANI* by a simple formula

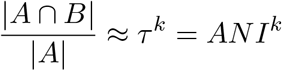

where the approximate equality can be made precise [63] under an appropriate random model.

The formula does not take into account coverage and read-sampling. To adjust for read sampling, we derive a different ANI formula by the following argument. We can write 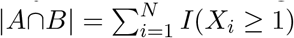 where *I* is the indicator 0-1 random variable. Thus, the expected value is

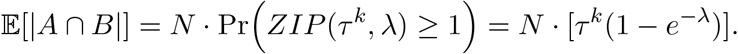

Remembering that we assume a fixed, non-random hash function for FracMinHash, so |*A*| = *N*, we get

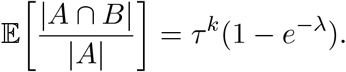

This immediately gives rise to our final ANI estimator, the *λ*−adjusted containment ANI estimator:

#### Definition 5.

*The coverage-adjusted ANI estimate is*

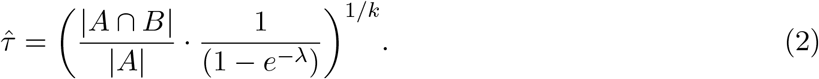

We can use this estimator once we know *λ*. In practice, we use the 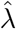 estimate in Equation 1 to first estimate *λ*, since this estimate does not depend on *τ*, and then we plug in the estimate 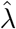 into Equation 2 to get our final ANI estimate 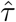. Note the *λ*−adjusted ANI in Equation 2 could be *>* 1, in which case we threshold *τ* = 1.

### 4.6 Practical thresholds

In practice, we apply three filters: (1) we only apply the *λ*−adjustment for ANI if the median multiplicity for k-mers *X*_1_, … is ≤ 3, as the term in Equation 2 is small if *λ* ≥ 3. (2) we only proceed with *λ*-adjustment if *N*_*a*_ ≥ 3 and *N*_*a*+1_ ≥ 3 where *a* is the k-mer multiplicity ≥ 1 with the most number of k-mers; this is usually *a* = 1. These filters are to make sure we only proceed with *λ* adjustment when it is necessary and if enough information is given. (3) we only output ANIs if the number of FracMinHash k-mers for a genome is *>* 50 by default, although this can be changed for smaller genomes and contigs.

### 4.7 Masking multi-copy and dependent k-mers

Note that we can remove or mask k-mers in *A*, the k-mers in our reference genomes, without affecting the inference of our parameters, even if that k-mer is in *B*. This masking procedure simply reduces our sample size during inference by discarding some of the sample *X*_*i*_s. We can thus remove FracMinHash k-mers that are not “satisfactory”. First, we only take unique FracMinHash k-mers in the reference genome, as our k-mer Poisson model is predicated on the assumption of unique k-mers. Second, we remove FracMinHash k-mers in the reference genome if they are less than 30 bp apart by default, as k-mers that are too close are strongly correlated, which is not suitable under our assumption of independence. We found the latter step gives better inferences for small *c*.

### 4.8 Deduplicating reads by locality-sensitive hashing and handling short paired-end fragments

For real short and paired-end reads, we found two major failures of our uniformly distributed reads assumption, which in turn impacts our Poisson coverage model. (1) For paired-end reads, if two reads for a pair overlap too much due to a small fragment length [64], k-mers can be double-counted. (2) PCR duplicates [65], such as from Illumina sequencing, can double-count reads and thus k-mers. To deal with (1), we use a simple heuristic: if a FracMinHash k-mer appears twice across a pair of reads, we consider this k-mer as counted only once. To deal with (2), we use a simple locality-sensitive hashing technique for deduplicating reads that we will describe for the rest of the section. In **Supplementary Figs. 16-18** we show that deduplication greatly improves detection for low-abundance genomes.

We first explain the deduplication algorithm for paired-end reads. We start by scanning through each read pair, and for each FracMinHash k-mer *x* detected, we take the first 32-mer on both the forward read and the reverse read, and we mask both 32-mers in a 010101…01 pattern, ending up with two 16-mers that we denote *y*_1_, *z*_1_. We get another pair of two 16-mers by masking with a 1010…10 pattern instead, which we call *y*_2_, *z*_2_. We then insert the tuples (*x, y*_1_, *z*_1_) and (*x, y*_2_, *z*_2_) into a set membership data structure (discussed below). If one or both of these two tuples are already present, we ignore the FracMinHash k-mer and assume it’s due to a PCR duplicate.

The above algorithm is locality sensitive in that if two PCR duplicates differ by at most 1 substitution, it is guaranteed to not double-count the k-mers coming from a PCR duplicate, as one of the two masking patterns has unchanged bases. In practice, it can often tolerate more. For our set membership data structure, we use a scalable cuckoo filter [66,67] implementation with false positive rate *p* = 0.0001, which we found to decrease memory consumption while not affecting inference.

For single-end reads, we make a few changes. First, we ignore reads with length *>* 400 bp, as these are likely to be long reads. Secondly, instead of letting *z*_1_ and *z*_2_ be from the second read in the pair, we take *z*_1_ and *z*_2_ to be the k-mers starting from the middle of the read. Finally, we only deduplicate for the FracMinHash k-mers with multiplicity *<* 4, as PCR duplicates are indistinguishable from true reads for high-coverage, single-end read sets.

### 4.9 Bootstrapping for uncertainty quantification

We found the main source of uncertainty to be in the estimation of *λ* due to read-sampling when *λ* is very small, which propagates to uncertainty in *τ* . Other sources of uncertainty, such as due to FracMinHash sampling, are not very large when the ANI is greater than 90% for complete bacterial genomes, as shown experimentally in [63].

To quantify the uncertainty in ANI due to *λ* uncertainty, we perform a bootstrap on the resulting distribution of *X*_*i*_s (the k-mer multiplicities). Sylph resamples (with replacement) the k-mer multiplicities over 100 iterations and performs coverage-adjusted ANI calculations for each iteration. Sylph then takes the 5th and 95th percentile ANI estimates as the 90% confidence interval. Notably, it is possible for a resampling that *N*_*a*_ or *N*_*a*+1_ is 0, in which case we throw out the observation. We only proceed if *>* 50 of the resamples give an *N*_*a*_ and *N*_*a*+1_ that are non-zero. Note that our bootstrapping procedure is fundamentally different from the issue encountered in a related study [68], where bootstrapping causes issues when resampling already-sampled reads; we resample *k-mers from the reference genome*, not k-mers from reads.

We show the confidence interval coverage probabilities (“coverage” here referring to confidence interval coverage, not read-sampling coverage) in **Supplementary Table 1** for a synthetic test. In general, sylph’s confidence intervals are slightly tight, but the coverage probabilities are not less than 70% under simulations. Under real samples with more sources of uncertainty, we expect our intervals to be tight but still provide a rough quantification of uncertainty.

### 4.10 Computation of effective and true coverage

Let *m* be the median k-mer multiplicity for a given genome. Effective coverage for a genome is output as either the *λ* estimated from Equation 1 if *m* ≤ 3, a robust mean k-mer coverage when 4 ≤ *m* ≤ 15, or just *m* otherwise. The robust mean k-mer coverage is calculated by taking the mean k-mer multiplicity for k-mers with multiplicity less than *α* defined as follows: *α* is the smallest number such that for *Pois*(*m*), a Poisson random variable parameterized by *m*, Pr(*Pois*(*m*) *> α*) *<* 10^−10^. We found that this robust mean is more accurate than the median for small (effective) coverages since the median is necessarily an integer. However, for large coverages, we found the robust mean is unnecessary, so we use the median instead.

The effective coverage, *λ*, can be much smaller than the true coverage, *δ*, due to read errors and small read lengths (Definition 2). Users can toggle sylph to estimate the true coverage instead using the following method. To estimate *δ*, we need to know the *L* and 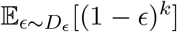, which we will denote *E*. This gives the resulting formula

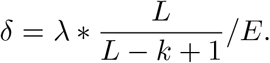

Sylph allows the user to input a fixed read length *L*. While read lengths for long reads may be heterogeneous, the effect of read length is negligible for long reads on the effective coverage, so this is not an issue in practice. For estimating *E*, if the user inputs a read error rate *ϵ*, sylph simply estimates *E* = (1 − *ϵ*)^*k*^. That is, *D*_*ϵ*_, the distribution of error rates, is a point distribution at *ϵ*. If the user does not input a read error rate, defining *n*_*x*_ as the number of k-mers in the read sketch with multiplicity *a*, sylph automatically estimates *E* as

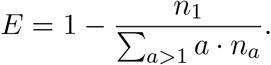

This formula comes from two assumptions: (1) lowly covered genomes are rare, so all k-mers with multiplicity 1 should be erroneous, and (2) erroneous k-mers are unique. Thus, ∑_*a>*1_ *an*_*a*_ · (1−*E*) ≈ *n*_1_, giving the above formula. In this study, we used sylph’s automatic *E* estimation for all samples not coming from soil or ocean samples. In soil and ocean samples, we found that assumption (1) does not hold and thus encourage users to provide an approximate *ϵ* of 0.005 to 0.01 for paired-end reads instead.

### 4.11 Profiling by k-mer reassignment

After ANI calculation, sylph can output results with the command *sylph query*. However, such results are not a metagenomic profile with accurate relative abundances. Shared k-mers within similar genomes in the database may cause multiple related species to have high ANI when only one of the species is actually present.

To reassign shared k-mers, we use a winner-take-all heuristic that is also an option in Mash screen and sourmash. The command *sylph profile* finds putative ANIs in the same manner as *sylph query*, but then assigns k-mers uniquely to the genome with the highest ANI. The ANI computation is done again but only considering these reassigned k-mers, giving our final ANIs. Sylph then outputs genomes for which ANI is *>* 95% by default.

### 4.12 Outputting abundances, percentage of reads detected, and taxonomic profiles

Let *GL*_1_, …, *GL*_*q*_ be the genome lengths of the *q* genomes passing the threshold. Let *λ*_*i*_ and *δ*_*i*_ denote the effective and true coverage of the *i*th detected genome. Sylph outputs the taxonomic abundance for each genome as 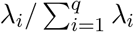. The sequence abundance output is 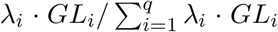. We estimate the percentage of reads detected at the species-level as 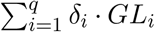 divided by the total number of bases in the reads times 100, outputting 100% if this value exceeds 100. Importantly, the percentage of reads detected depends on the *true* coverage.

We provide a script for turning a metagenomic profile without taxonomy information into a taxonomic profile for multiple pre-built databases. This script can be easily customized by adding taxonomic accessions to arbitrary genomes. This is done by translating each genome in the output to a taxonomic accession string and aggregating sylph’s output abundances at each taxonomic rank and is independent of the core algorithm.

### 4.13 Synthetic metagenome construction and benchmarking

To construct the undercharacterized synthetic metagenome (**Fig. 2A**), we first computed nearest neighbor ANIs from the newer GTDB-R214 database to GTDB-R89. All genome-to-genome ANI calculations were done with skani [69]. We then arbitrarily selected 50 genomes with 95-97.5% ANI and 150 genomes with 85-90% ANI (**Fig. 2A**) nearest neighbour ANI. We then sampled 3 Gbp of 2x150bp reads for this dataset using wgsim [70] with abundance following a log-normal distribution with mean and standard deviation of the underlying normal distribution equal to 2 and 1 respectively, as is done in other benchmarks [71]. 10 samples were created for this dataset. Exact software commands are shown in Supplementary Methods C. For the 5 synthetic metagenomes binned by 1% ANIs (**Fig. 2B**), for each 1% ANI bin, we generated a metagenome with 50 genomes each and 750M bp of reads using the same distributions as for the undercharacterized metagenome case.

MetaPhlAn4 could not be profiled against the undercharacterized case because it contains genomes in the holdout set. We attempted to profile MetaPhlAn4 on the 95-100% binned ANI dataset – its database should contain all GTDB-R89 species as its taxonomy can be mapped to GTDB-R207, a newer database [11]. Therefore, we first mapped taxa from MetaPhlAn4’s default NCBI taxonomy to the GTDB-R207 database as provided by a script included in MetaPhlAn4. Then, we projected the representative genome for the GTDB-R207 taxa to the nearest neighbour ANI genome’s species name in the GTDB-R89. An issue we found was that MetaPhlAn4 SGBs could collapse species together in the GTDB-R89 taxonomy, thus picking the wrong representative due to the inherent incompatibility with the SGB definition and GTDB. This lowers the precision and sensitivity (**Supplementary Figs. 4 and 9**).

### 4.14 CAMI2 benchmarking details

To obtain CAMI2 results, we took the profiles for MetaPhlAn2.9, mOTUsv2.5, and KMCP as compiled in KMCP’s study [30] and the official CAMI2 repository. We reran all profiles using the latest version of OPAL [33] (**Supplementary Table 2**) due to finding slightly different values depending on the version of OPAL. For sylph, we took all genomes in the provided RefSeq CAMI2 snapshot (Jan 8 2019) with a taxid present and used these genomes as sylph’s database (with default settings). TaxonKit [72] was used for converting sylph’s output into a CAMI-compatible profile.

We also ran MetaPhlAn4 and mOTUs3 ourselves due to no official profiles being available. However, these profilers use more comprehensive databases than the official submissions, so direct comparison is not completely fair. We discuss taxonomy harmonization for CAMI2 in Supplementary Methods B.

In the Marine dataset, we found that for the detected species, many had closely related genomes in the provided database; sylph’s median ANI was 99.8% over all samples for detected genomes in the Marine dataset. There are also species in the true profiles that are not present in the given reference genomes, obfuscating the profiling metrics. For example, Psychrobacter sp. JCM 18900 is present in the taxonomy and in the gold standard profile for multiple samples in the Marine dataset, but it is not present in the RefSeq databases and is not found by any method. This causes other Psychrobacter species to be profiled and classified as incorrect, lowering the sensitivity, because the missing genome can not be detected, but *also the precision* because “incorrect” but related species will be identified instead.

### 4.15 MWAS procedure details

We ran logistic regression using statsmodels [73] for each of the 289,232 genomes with sylph’s ANI as a covariate and the disease-control status as the response. In addition, we used all of the same metadata covariates as specified by Table 1 in Wallen et al. [38] to control for potential confounding (e.g. sex, age). Note that we only used one genome per species representative in the Q-Q plot (**Fig. 4D**) because genomes in the same species give highly dependent p-values, but we ran the MWAS on all 289,232 genomes.

We set a presence-absence threshold of 98% ANI, meaning if the ANI for a sample was below 98%, we set it to 98% for the regression. Otherwise, if the ANI is *>* 98%, we keep it as is. This gives a continuous presence-absence covariate so that only high ANIs give information for the regression, which is desirable when trying to find strain-level associations. We then used the Benjamini-Hochberg procedure [74] to control for FDR at a significance level of *α* = 0.05.

We created a linear ordering on genomes within a species group, where similar genomes should be placed next to each other along the Manhattan plot, by running an ANI-based average-linkage clustering on the genomes. We used skani [69] for all-to-all ANI calculation and scipy [75] for clustering, using the resulting dendrogram ordering as the within-species ordering.

### 4.16 Synthetic viral metagenome profiling

To generate synthetic viral metagenomes for profiling, we downloaded all viral genomes from RefSeq and the MGV database [48] and computed ANI from all RefSeq viruses to MGV using skani and the <monospace>--slow</monospace> setting. RefSeq viral genomes with *>* 95% ANI and *>* 80% aligned fraction for both the query and the genome were considered, with the nearest neighbor MGV genome being designed as the true genome. We randomly selected 50 genomes for each sample over 10 samples and simulated reads using the same methodology as with the other synthetic metagenomes, except we only sampled 3Mbp of reads for each sample.

### 4.17 Benchmarking hardware and software

Benchmarking was done on an Intel(R) Xeon(R) CPU @ 3.10GHz machine with 64 cores and 240 GB of RAM as a Google Cloud instance with an SSD disk. Software versions are shown in **Supplementary Table 2**.

## 5 Data availability

GTDB databases can be found at https://gtdb.ecogenomic.org/. CAMI2 datasets were taken from https://data.cami-challenge.org/participate, with KMCP profiling results taken from https://doi.org/10.5281/zenodo.7450803. Meslier et al. raw data and metadata are available at PRJEB52977 and additional scripts and references can be found at https://forgemia.inra.fr/metagenopolis/benchmark_mock. Wallen et al. raw data and metadata are available at PR-JNA834801. Chng et al. read data is available at PRJNA277905 and patient metadata is available in Supplementary Table 4 of the study by Chng et al [47]. The MGV and IMGVR databases are available at https://portal.nersc.gov/MGV/ and https://genome.jgi.doe.gov/portal/IMG_VR/IMG_VR.home.html. Mouse gut metagenomes are publicly available with accession PR-JNA549182. Biofloc metagenomes are publicly available with accession PRJNA967453, and the associated MAGs are available at https://doi.org/10.6084/m9.figshare.23599461.50 real gut metagenomes used for benchmarking are available in Supplementary Table 4.

## 6 Code availability

sylph is available at https://github.com/bluenote-1577/sylph. Data analysis scripts and notebooks can be found at https://github.com/bluenote-1577/sylph-test.

## 7 Competing interest statement

No competing interests declared.

## 8 Acknowledgements

J.S. was supported by an NSERC CGS-D scholarship. This work was supported by Natural Sciences and Engineering Research Council of Canada (NSERC) grant RGPIN-2022-03074 and DND/NSERC Supplement DGDND-2022-03074.

## A MOCK2 dataset from Meslier et al. [34]

We took a previously published synthetic mock community with 87 known diverse microbial genomes (MOCK2 community from Meslier et al. [34]) sequenced with real PacBio HiFi, Oxford Nanopore R9, and Illumina HiSeq3000 technologies, shown in Fig. 1C. Notably, the PacBio HiFi sequencing run appears to be mislabeled in the study; the authors claim the PacBio run was sequenced from the MOCK1 community but we detected various species not present in MOCK2. For example, *Streptococcus agalactiae* was claimed to be only present in MOCK2, but the PacBio run for MOCK1 had this species detected by sylph at 100% ANI, indicating that MOCK2 was actually sequenced instead.

## B CAMI2 profiling with MetaPhlAn4 and mOTUs3

The usage of MetaPhlAn4 and mOTUs3 profiles in our results for the CAMI2 benchmarking were not official submissions for the CAMI2 challenge – to our knowledge, no official submissions for MetaPhlAn4 or mOTUs3 that are concordant with the official CAMI2 databases exist. As such, we reran MetaPhlAn4 and mOTUs3 (in precision mode) using their respective options for outputting results in CAMI format.

Both MetaPhlAn4 and mOTUs3 may contain more genomes in their databases than the standard databases used for the official CAMI2 submissions, so evaluation is not necessarily fair. Furthermore, note that CAMI2 uses the Jan 8, 2019 NCBI taxonomy. While mOTUs3 adheres to this taxonomy [12], MetaPhlAn4 may not (https://forum.biobakery.org/t/which-ncbi-taxdump-version-used-for-metaphlan4-database/4989).

## C Profiling commands and information

We used the GTDB-R89 dereplicated genome database for Fig. 2A,B. We converted the GTDB taxonomy to the taxonomy dump files using https://github.com/rrwick/Metagenomics-Index-Correction [76]. We then used each methods method for building their custom database using this taxonomy.

### C.1 sylph

~~~
sylph sketch -1 A_1.fq B_2.fq … -2 A_2.fq B_2.fq … -d sylph_sketches -t 50 sylph profile sylph_sketches/* my_databases.syldb -t 50 > output.tsv
~~~

### C.2 Kraken2 + Bracken (0.01% sequence abundance cutoff used)

~~~
kraken2 --paired --db database --threads 50 r1.fq r2.fq
\\ --report kraken_out/sample.kreport;
bracken -d database -i kraken_out/sample.kreport -o output -r 150
~~~

### C.3 KMCP

~~~
kmcp search -d db_folder/db.kmcp -o sample.kmcp.tsv.gz -1 r1.fq -2 r2.fq -j 50 kmcp profile -X db_folder/ -T db_folder/taxid.map -m 3
\\ sample.kmcp.tsv.gz -o output -C output.profile -s sample_name
~~~

### C.4 ganon

~~~
ganon classify -d ganon_db -p r1.fq r2.fq -o output -t 50
~~~

### C.5 MetaPhlAn4

~~~
metaphlan r1.fq,r2.fq --bowtie2out metaphlan_out/sample.bowtie2.bz2
\\--nproc 50 --input_type fastq -o output
~~~

### C.6 mOTUs v3

~~~
motus profile -f r1.fq -r r2.fq -t 50 -A -o output
~~~

**Supplementary Figure 1:**
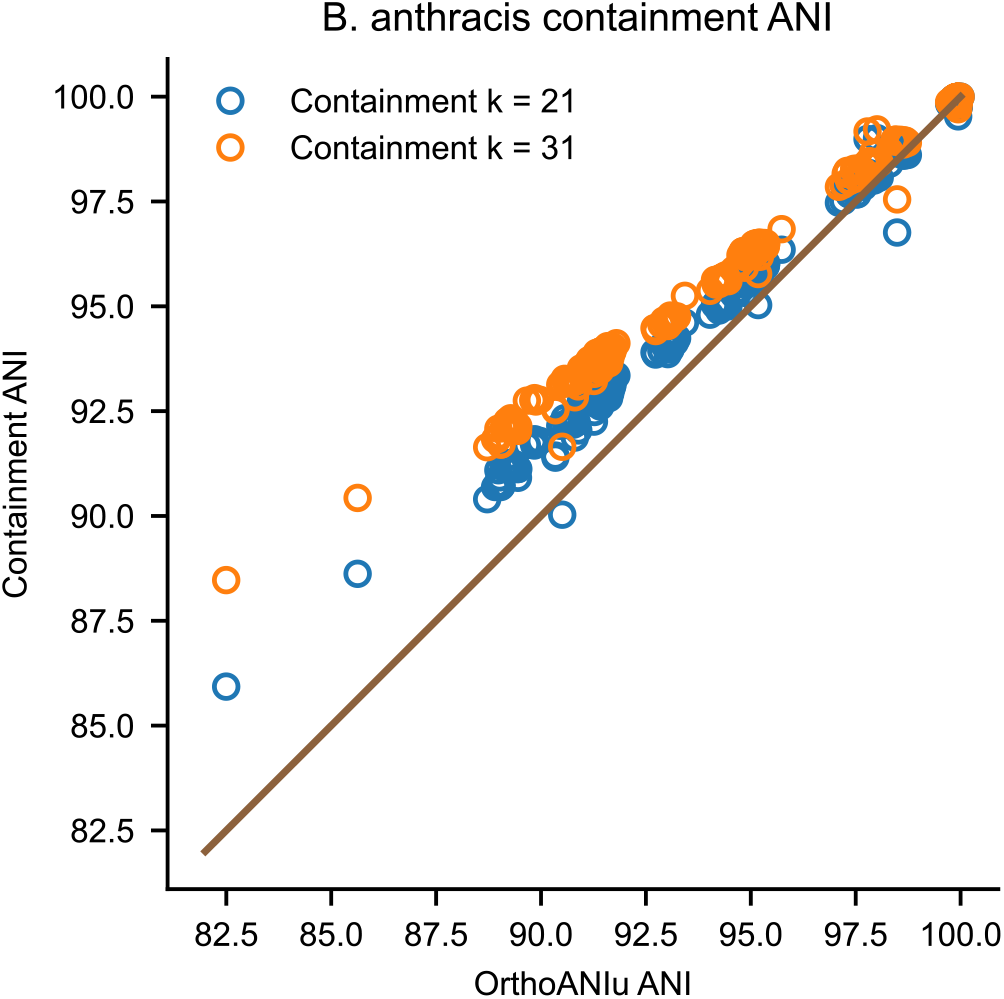
Containment ANI compared to OrthoANI [77] for a *bacillus anthracis* genome against a database of *B. anthracis* genomes taken dataset D2 from Jain et al. [27]. k=21 or k=31 indicates the k-mer size.

**Supplementary Table 1:** Coverage probabilities of sylph’s estimated 90% confidence intervals in Supp Fig. 2 if sylph outputs an interval, i.e. 100 times the fraction that sylph’s 90% confidence interval covered the true ANI over all data points (n = 200). Sylph only outputs a confidence interval when there are enough shared k-mers.

| Reference genome (ANI with *K. pneumoniae*) | $c = 100$ | $c = 200$ | $c = 1000$ |
| --- | --- | --- | --- |
| *K. pneumoniae* (100% containment ANI) | 78 | 72 | 87. |
| *K. africana* (96.2% containment ANI) | 70 | 84 | 91 |
| *K. aerogenes* (90.5% containment ANI) | 83 | 88 | 82 |

**Supplementary Figure 2:**
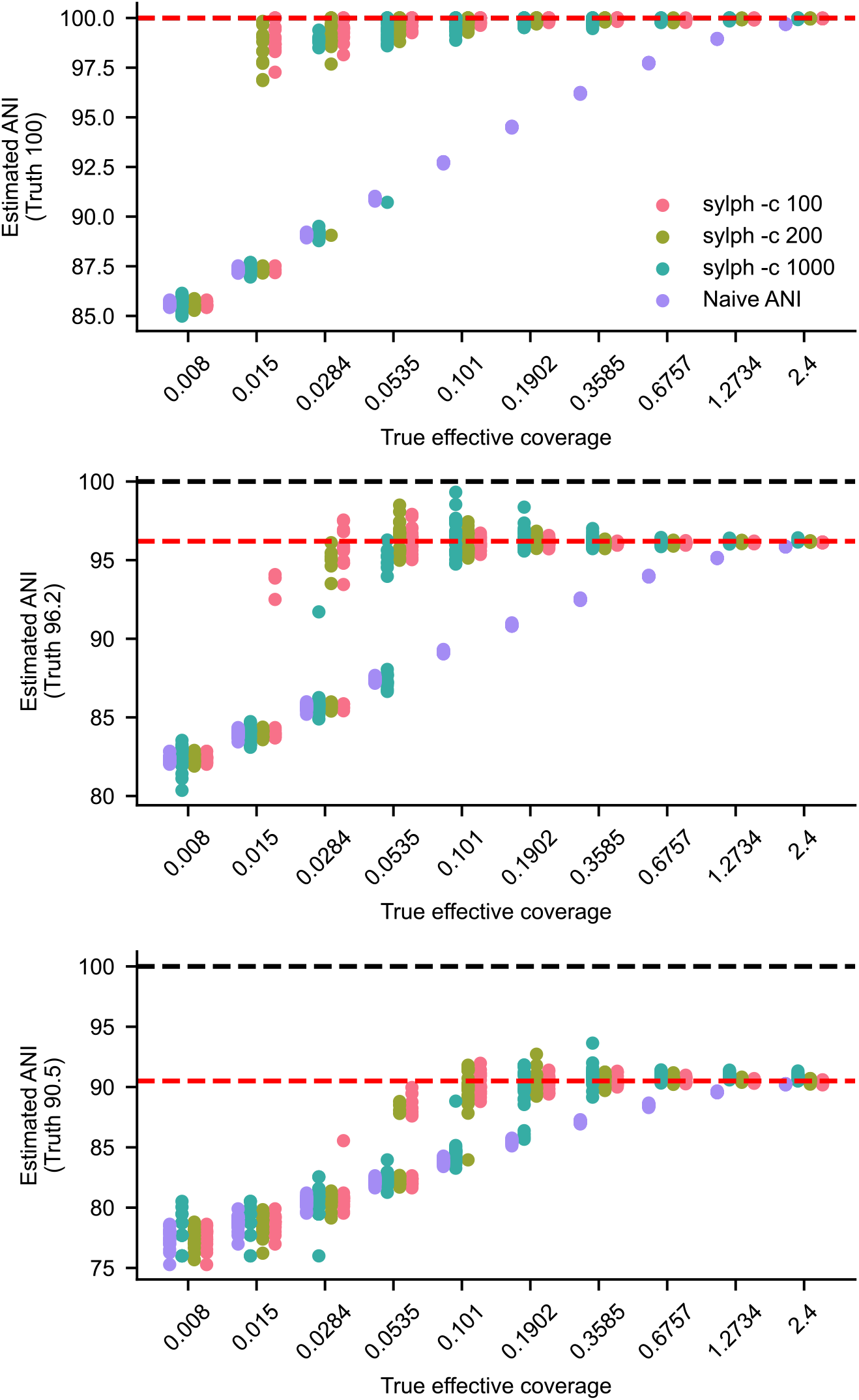
Naive containment ANI versus sylph’s adjusted ANI estimates when a genome is queried against a set of down-sampled, error-free simulated reads. We simulated 20 sets of reads from *K. pneumoniae* and queried *K. pneumoniae, K. africana*, and *K. aerogenes* against the reads from top to bottom subfigures. The true ANI is the true 31-mer containment ANI between *K. pneumoniae* and the respective genome. The coverage adjustment is crucial at lower coverages, and it works better when the true ANI is large and *c*, the subsampling rate, is smaller. Naive containment ANI calculated with sylph and *c* = 1000. The True ANI is calculated as the 31-mer containment ANI between the two genomes.

**Supplementary Figure 3:**
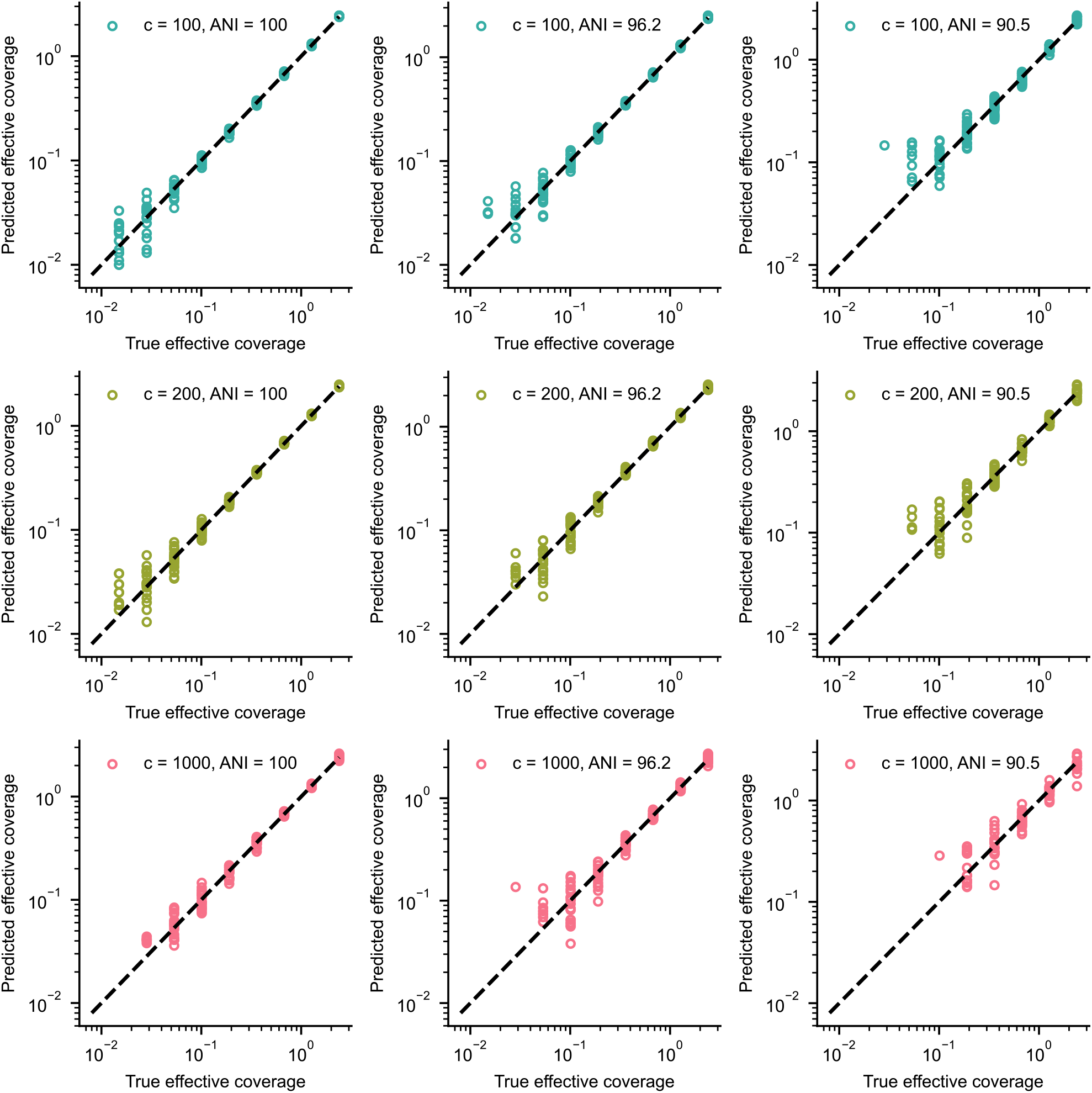
sylph’s *λ* (effective coverage) estimator versus the true effective coverage on the same synthetic *Klebsiella* dataset as from Fig. 2. Points were not shown if sylph could not output an estimate for *λ*.

**Supplementary Figure 4:**
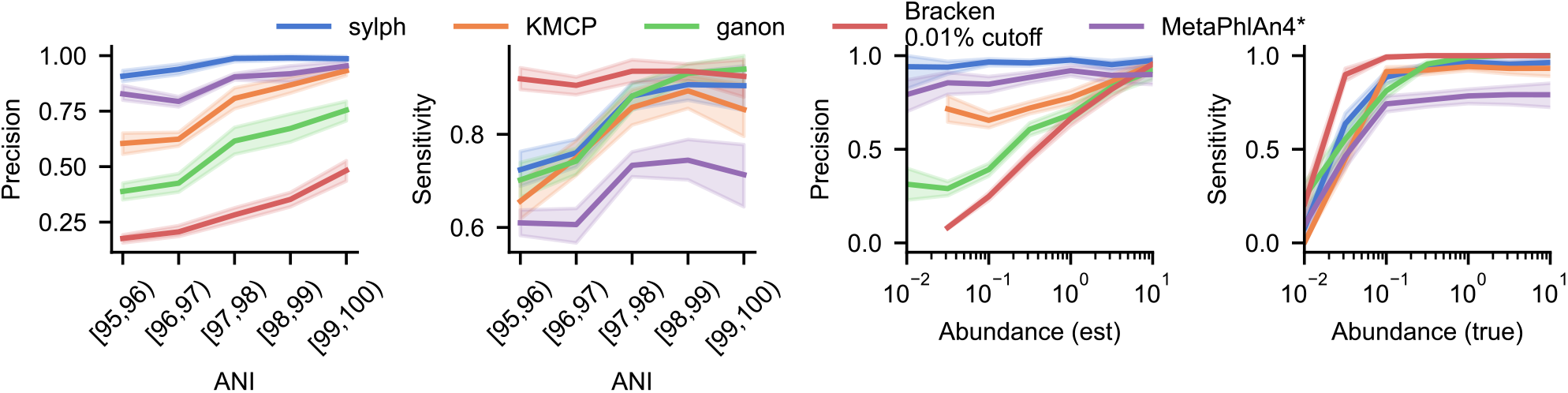
MetaPhlAn4 profiling on the combined synthetic metagenomes with 95-100% ANI to the GTDB-R89 database (**Fig. 2B**) with taxonomy harmonization as described in Methods.

**Supplementary Figure 5:**
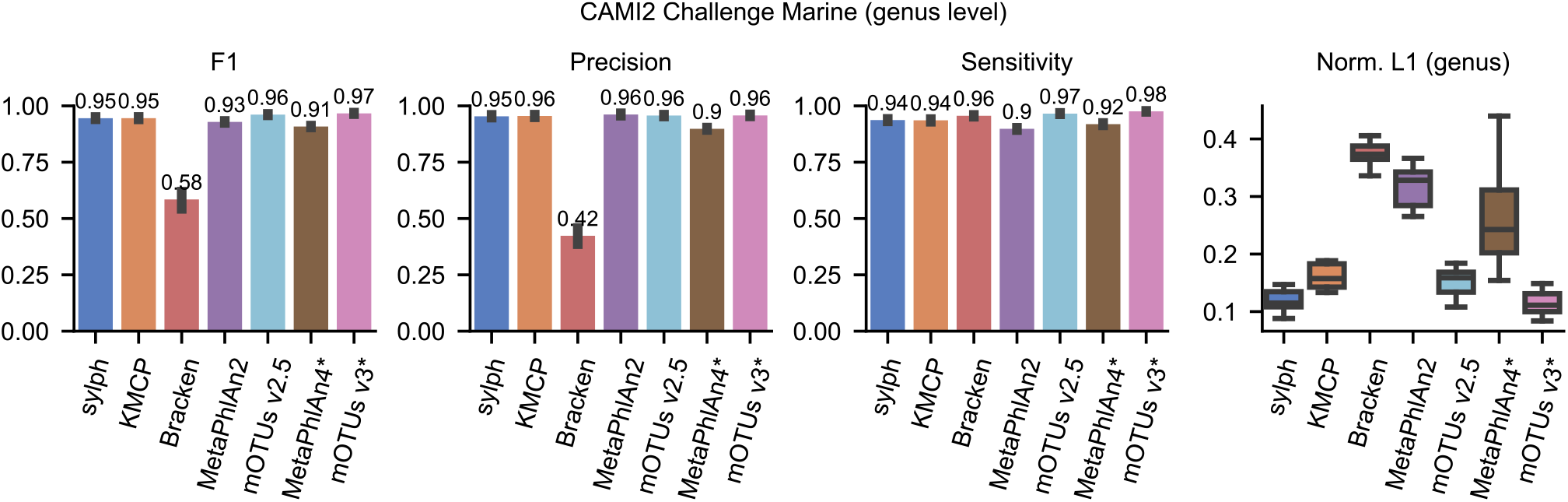
Profiling results for the CAMI2 Challenge Marine dataset at the genus level instead of at the species level, including methods that do not use CAMI2’s official RefSeq database or taxonomy snapshot (marked with an asterisk).

**Supplementary Figure 6:**
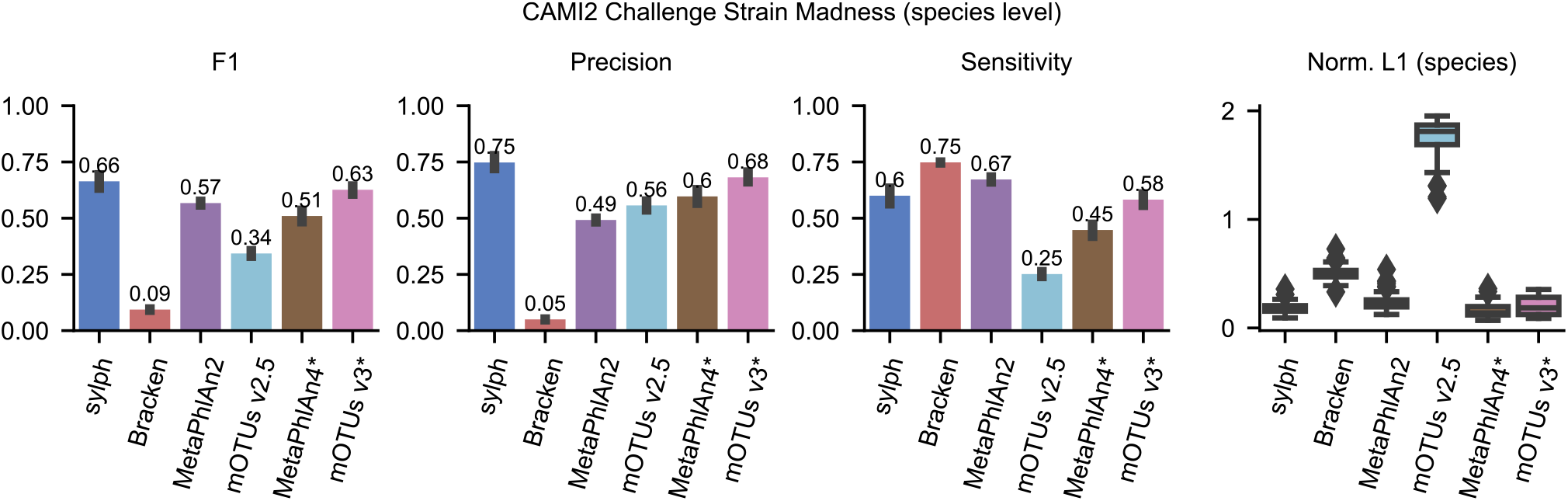
Profiling results for the CAMI2 Challenge Strain Madness dataset at the species level, including methods that do not use CAMI2’s official RefSeq database or taxonomy snapshot (marked with an asterisk).

**Supplementary Figure 7:**
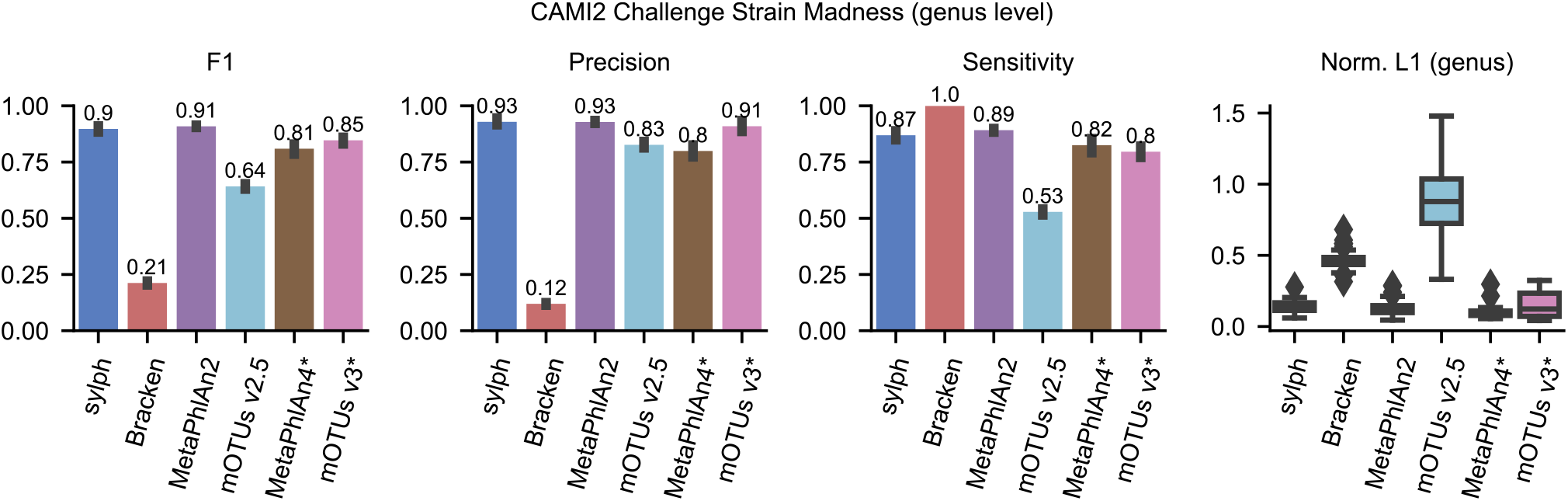
Profiling results for the CAMI2 Strain Madness dataset at the genus level instead of at the species level, including methods that do not use CAMI2’s official RefSeq database or taxonomy snapshot (marked with an asterisk).

**Supplementary Figure 8:**
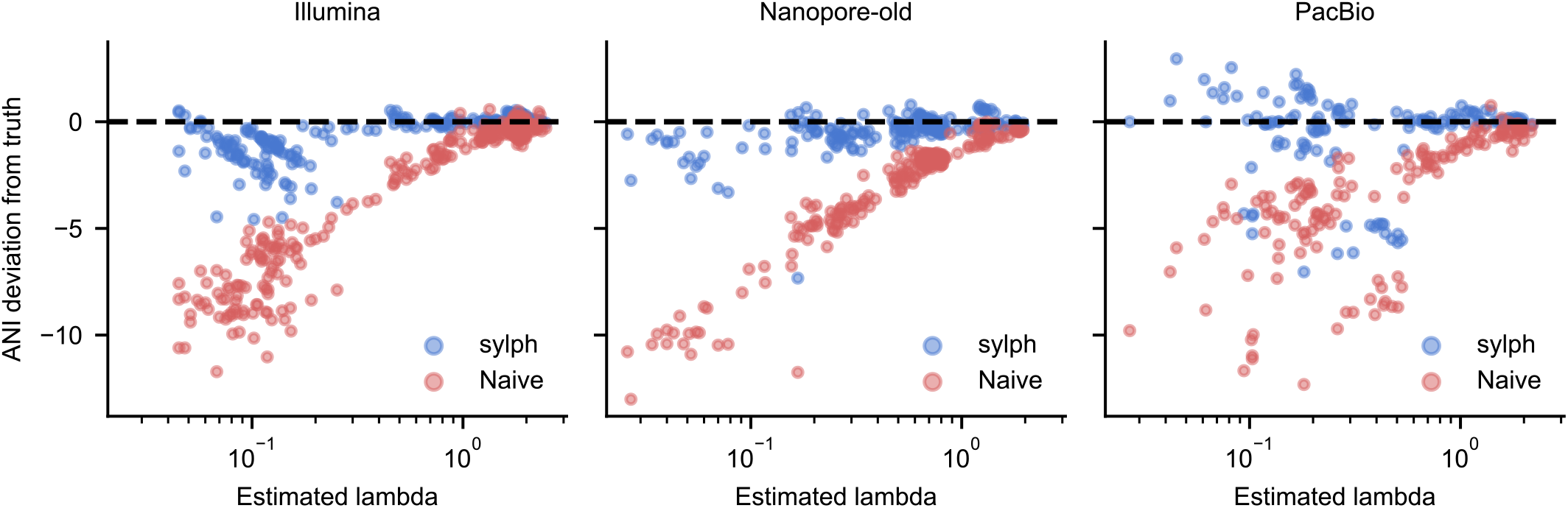
Downsampled Meslier et al. [34] dataset queried against the GTDB database, except ‘s ANIs are plotted as deviations from the true nearest neighbour ANI as a function of the estimated effective coverage parameter (*λ*) as estimated by sylph.

**Supplementary Figure 9:**
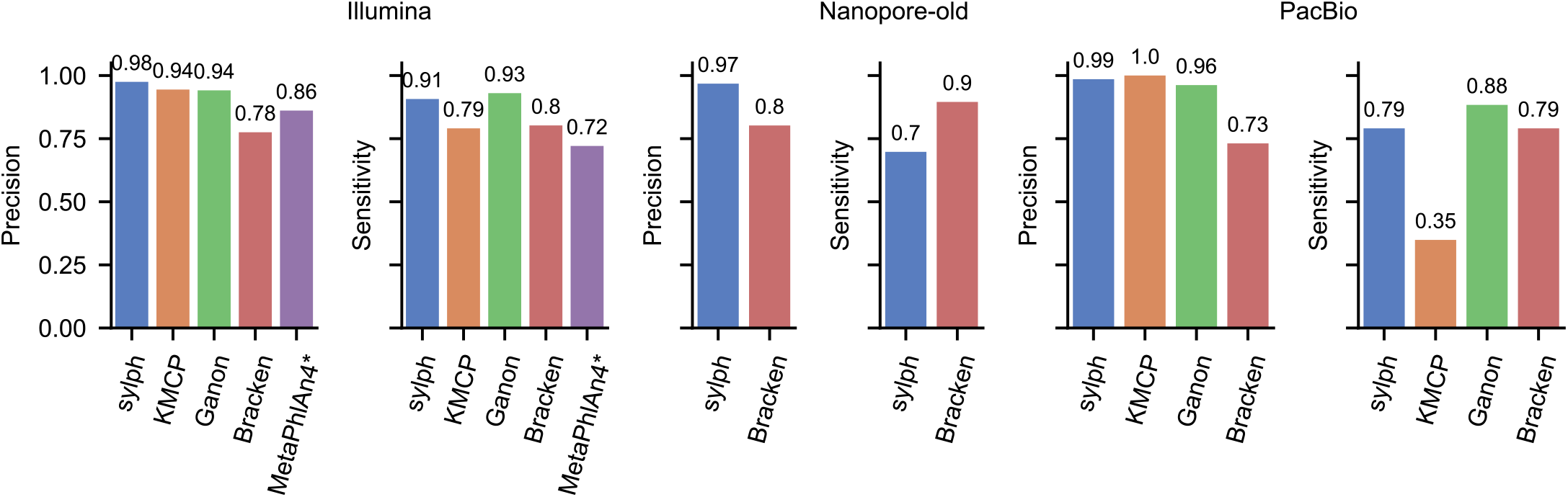
Profiling with on the downsampled Meslier et al. database (**Fig. 3C**) but with MetaPhlAn4 included. GTDB-R89 taxonomy harmonization with MetaPhlAn4 is described in Methods.

**Supplementary Figure 10:**
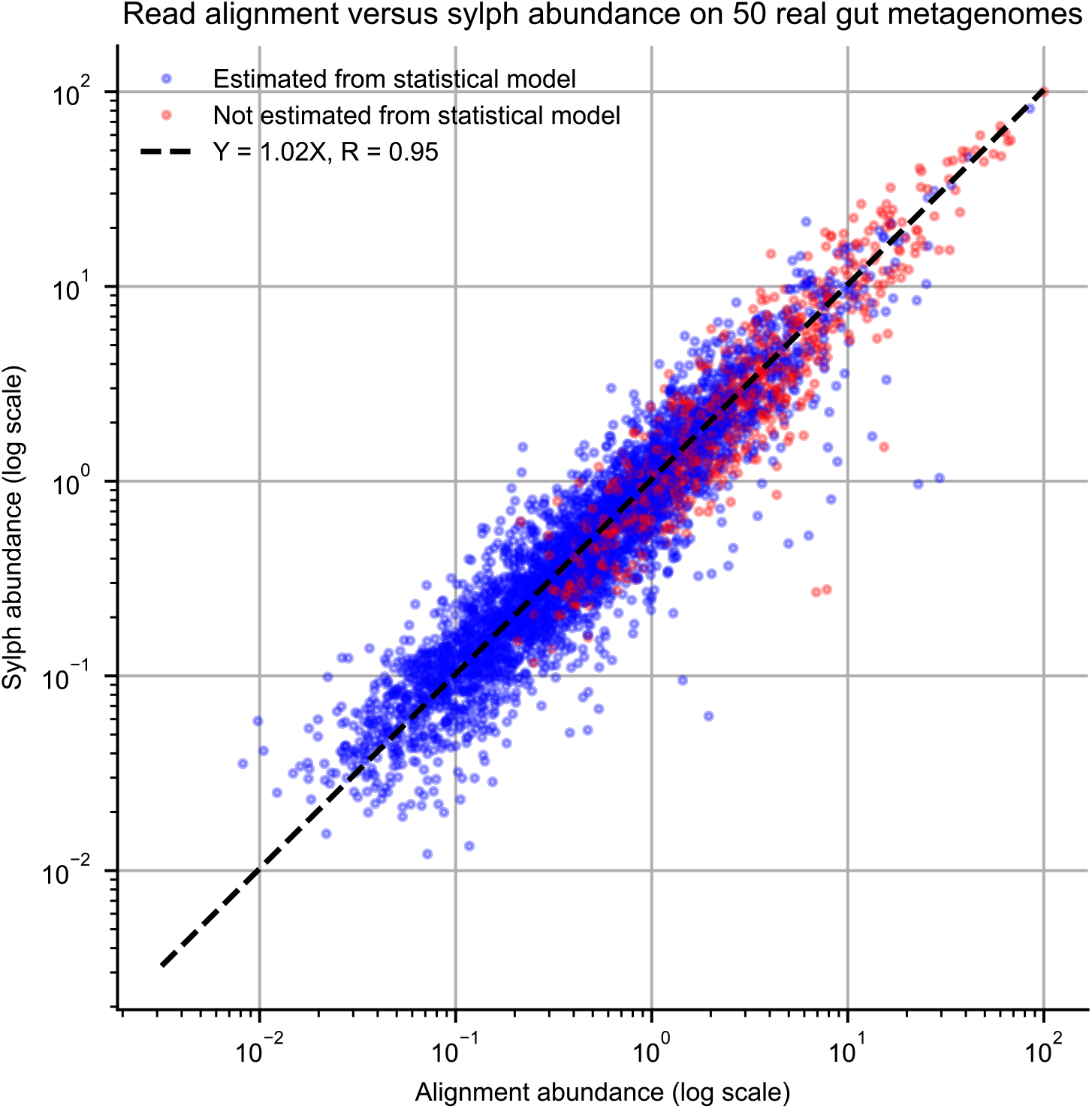
Profiling with 50 randomly selected short-read gut metagenomes from GMrepo v2 [36] with sylph and re-aligning the reads to sylph’s putatively present genomes to calculate abundances. Coverage was calculated with CoverM [78] and reads were aligned with minimap2 [79]. Red dots are genomes with coverages estimated by our zero-inflated Poisson model whereas blue dots are estimated using medians or means of k-mer multiplicities (see **Methods**).

**Supplementary Figure 11:**
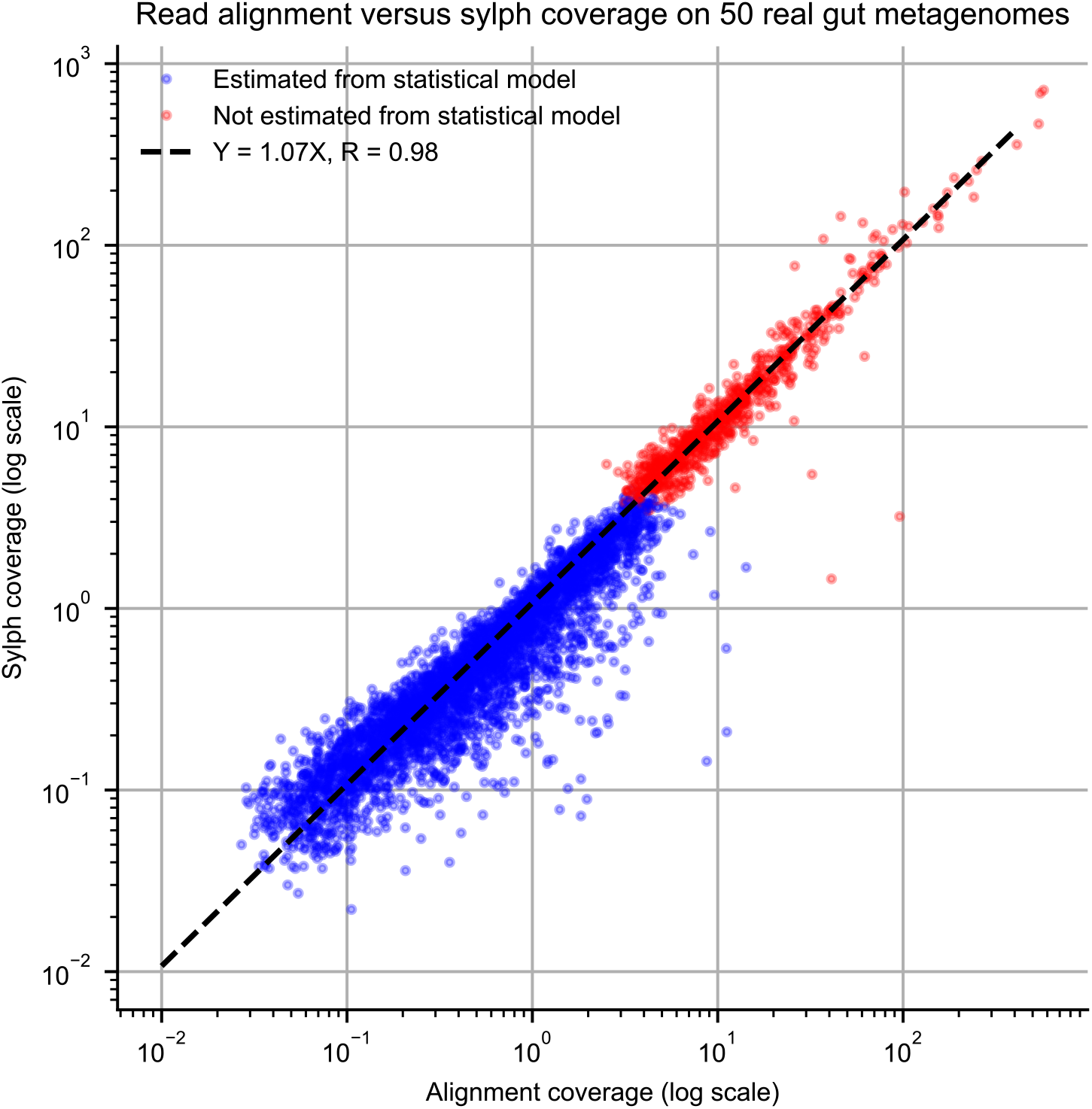
The same experiment and dataset as in Supplementary Figure 10 but with coverage instead of abundance shown. Coverage was estimated by arbitrarily assuming a sequence identity of 99.7% across all datasets, corresponding to 0.3% error rate for Illumina reads [80].

**Supplementary Figure 12:**
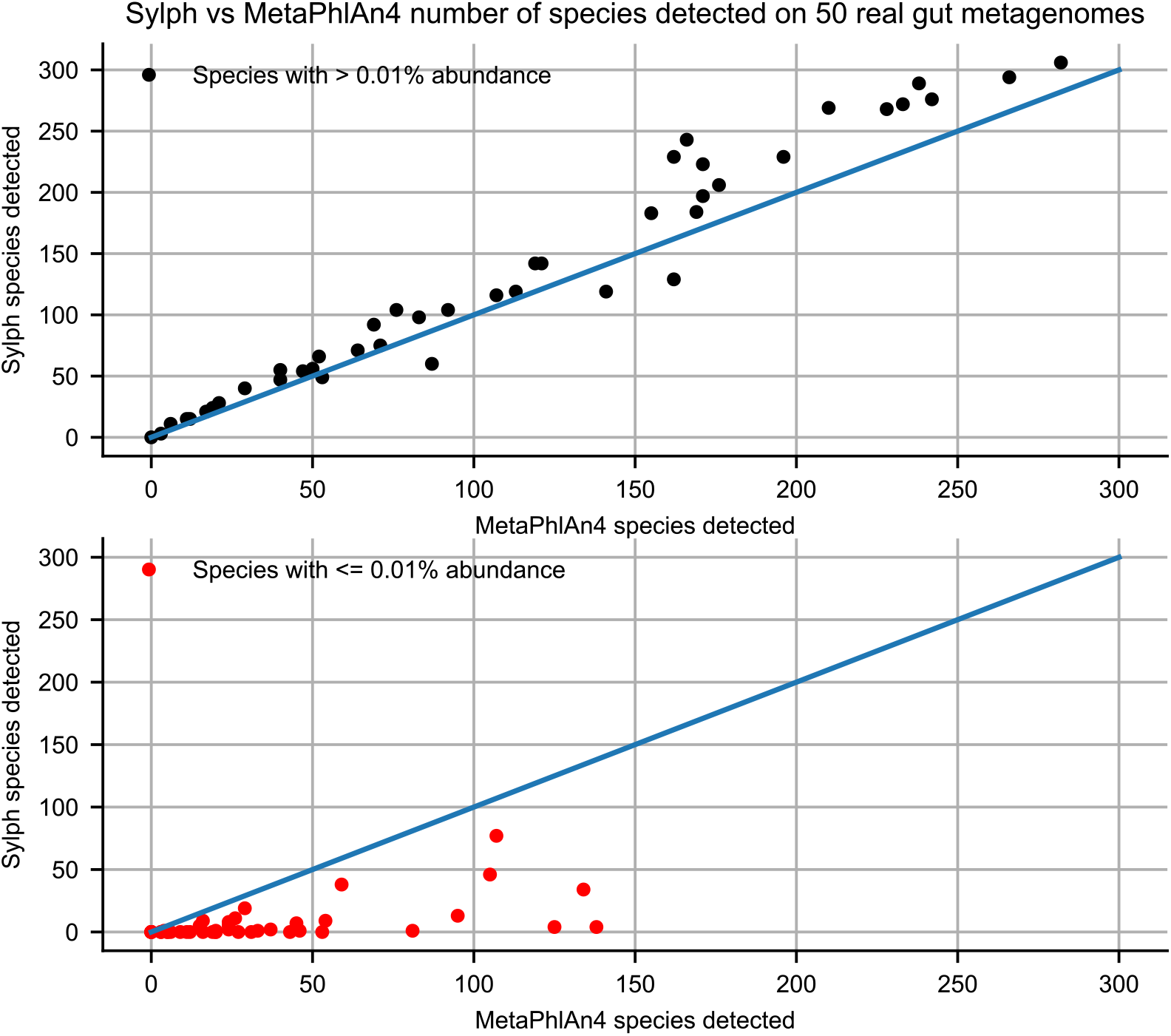
Plotting sylph’s number of predicted species versus MetaPhlAn4 for species with *>* 0.01% abundance (top) and ≤ 0.01% abundance (bottom). The same 50 gut metagenome dataset as in Supplementary Figure 10 was used. Both profilers were converted to the GTDB-R207 taxonomy.

**Supplementary Figure 13:**
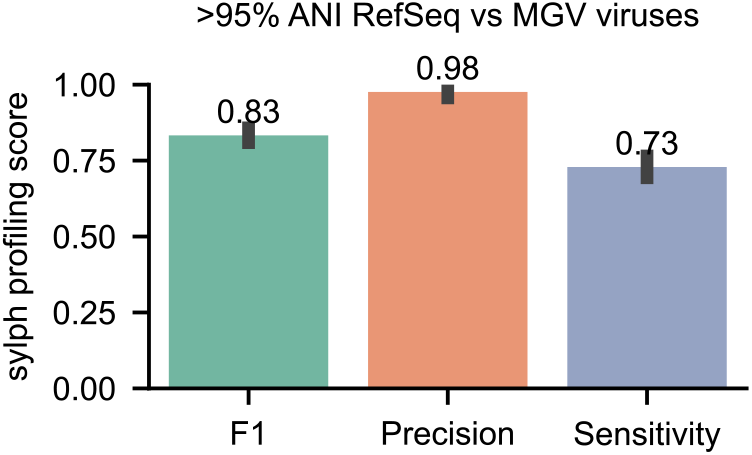
Sylph’s profiling results for synthetic communities of RefSeq viral genomes against a database of MGV viral genomes [48] with 95% confidence intervals shown over 10 samples.

**Supplementary Figure 14:**
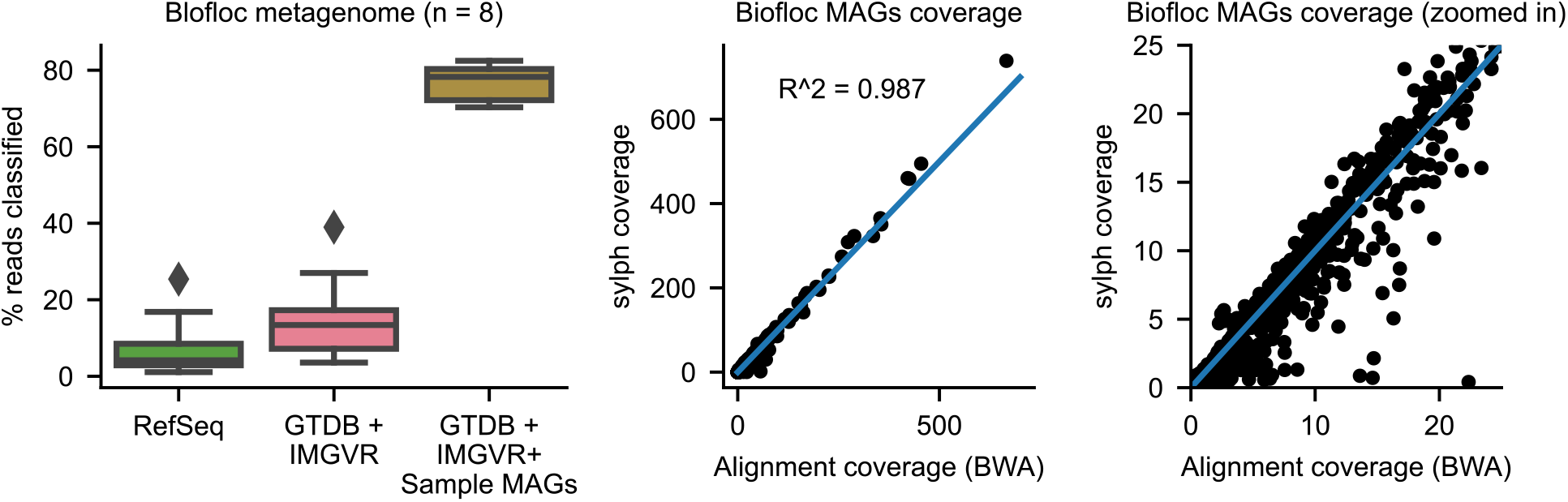
Coverage estimates output by sylph (y-axis) for the biofloc MAGs versus coverages obtained from read alignment using BWA (x-axis) with Pearson correlation coefficient shown.

**Supplementary Figure 15:**
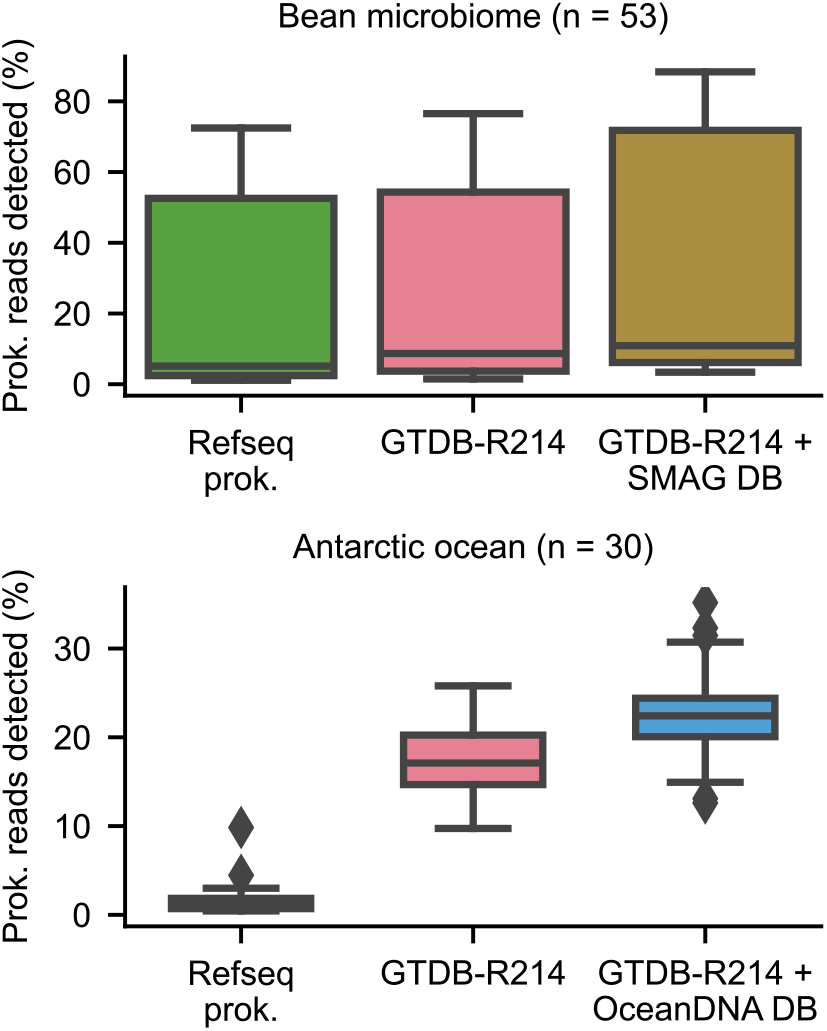
Profiling of plant-associated metagenomes [81] (PRJNA904562) and ocean metagenomes [82] (PRJEB61010) against ocean MAGs (OceanDNA) [52] and Soil MAGs (SMAG) [53]. We checked that these metagenomes were not included in the generation of the databases.

**Supplementary Table 2:**
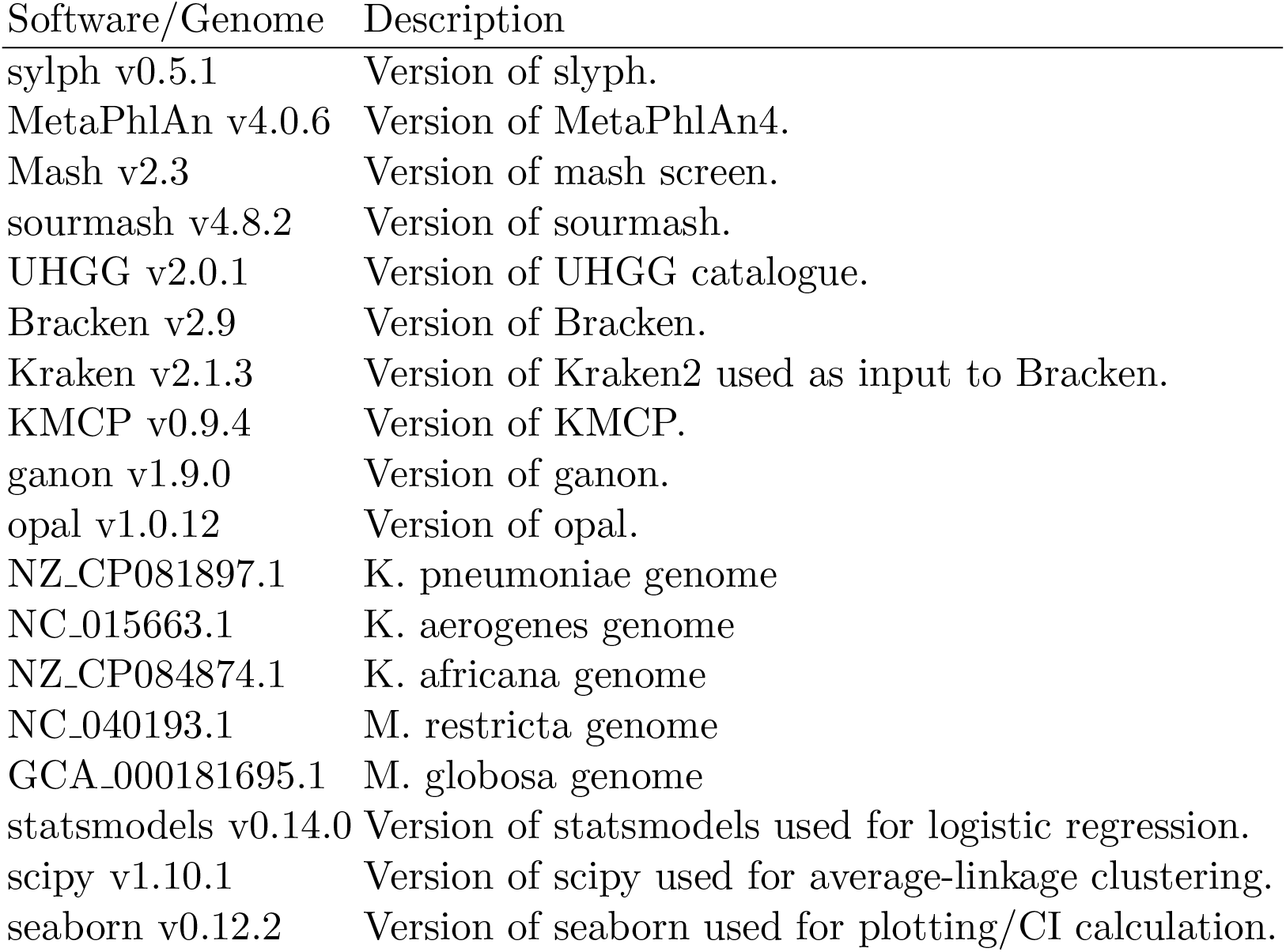
Software, genome, and data usage information.

**Supplementary Figure 16:**
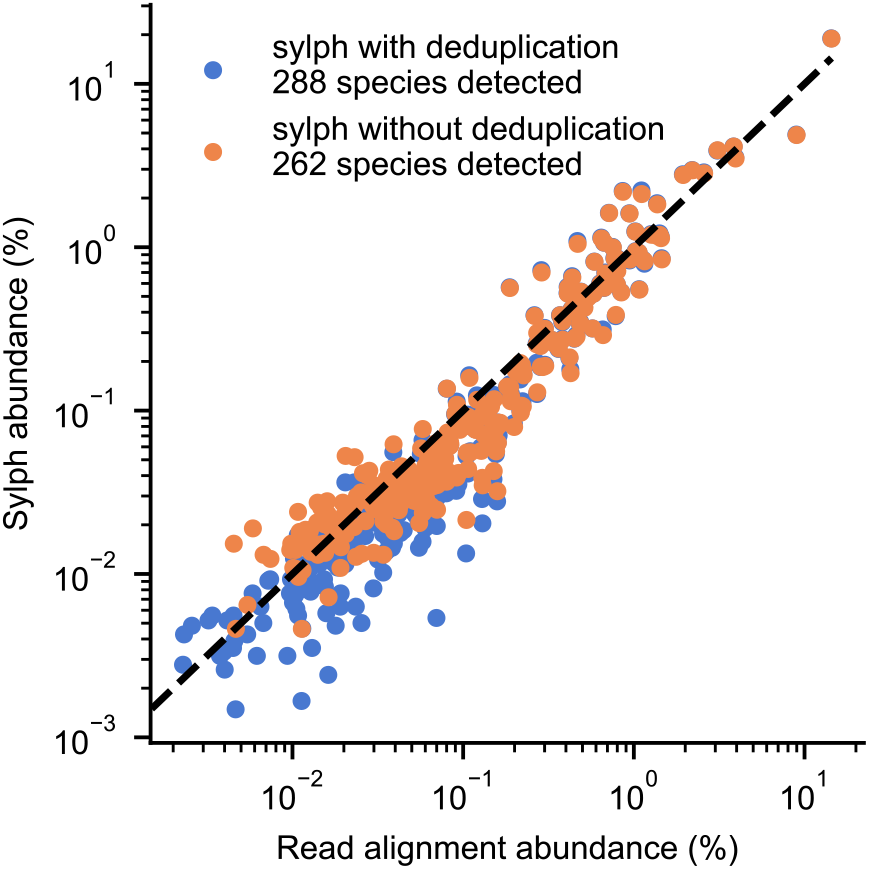
Sylph versus read mapping for before and after PCR deduplication for reads (SRR5983472) with 6.33% deduplication rate. Coverage and abundance were calculated using CoverM for the genomes that were present in the deduplicated results.

**Supplementary Figure 17:**
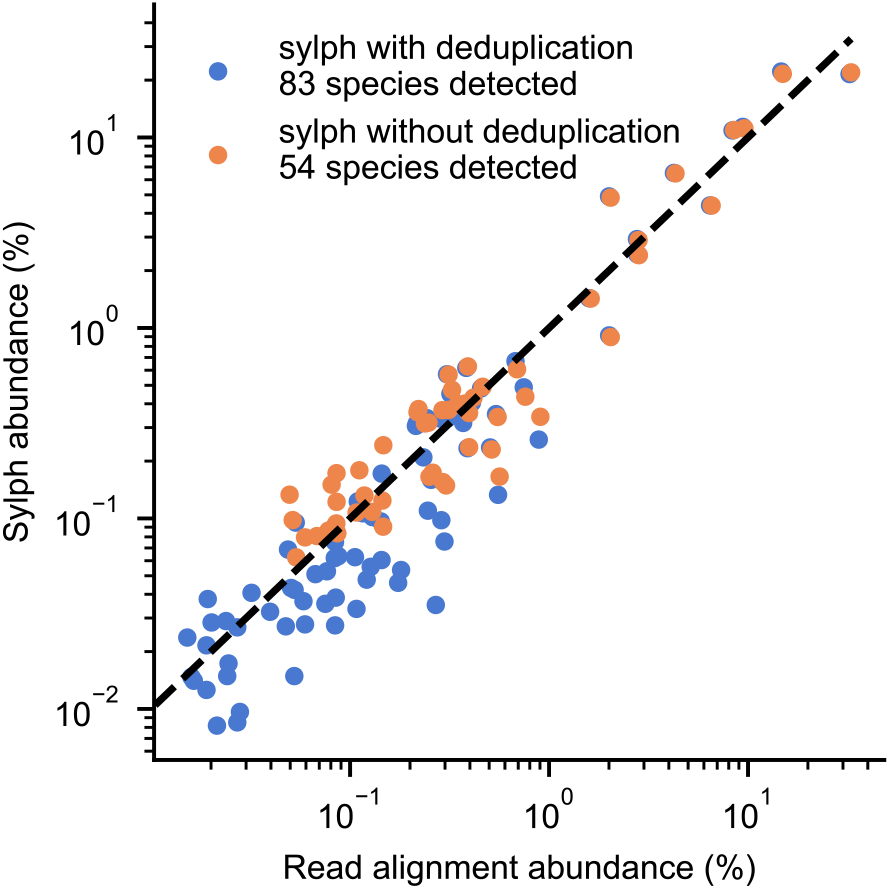
Sylph versus read mapping for before and after PCR deduplication for reads (SRR6075232) with 10.94% deduplication rate.

**Supplementary Figure 18:**
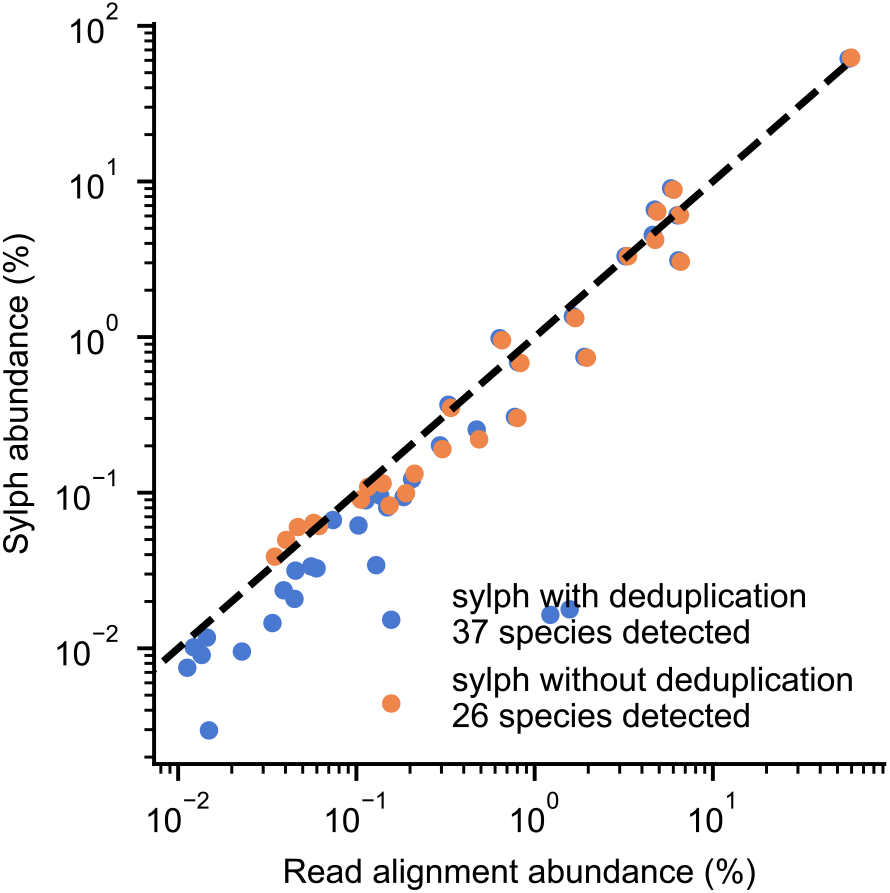
Sylph versus read mapping for before and after PCR deduplication for reads (SRR5983313) with 15.91% deduplication rate.

## Notes

https://github.com/bluenote-1577/sylph

https://github.com/bluenote-1577/sylph-test

### Competing Interest Statement

The authors have declared no competing interest.

### Summary of Updates

The results have been updated to include additional synthetic and real experiments. The methods have been updated with respect to sylph v0.5.1, including a deduplication hashing procedure.

